# Neuronal brain region-specific DNA methylation and chromatin accessibility are associated with neuropsychiatric disease heritability

**DOI:** 10.1101/120386

**Authors:** Lindsay F. Rizzardi, Peter F. Hickey, Varenka Rodriguez DiBlasi, Rakel Tryggvadóttir, Colin M. Callahan, Adrian Idrizi, Kasper D. Hansen, Andrew P. Feinberg

## Abstract

Epigenetic modifications confer stable transcriptional patterns in the brain, and both normal and abnormal brain function involve specialized brain regions, yet little is known about brain region-specific epigenetic differences. Here, we compared prefrontal cortex, anterior cingulate gyrus, hippocampus and nucleus accumbens from 6 individuals, performing whole genome bisulfite sequencing for DNA methylation. In addition, we have performed ATAC-seq for chromatin accessibility, and RNA-seq for gene expression in the nucleus accumbens and prefrontal cortex from 6 additional individuals. We found substantial neuron- and brain region-specific differences in both DNA methylation and chromatin accessibility which were largely non-overlapping, and were greatest between nucleus accumbens and the other regions. In contrast, glial methylation and chromatin were relatively homogeneous across brain regions, although neuron/glia ratios varied greatly, demonstrating the necessity for cellular fractionation. Gene expression was also largely the same across glia from different brain regions and substantially different for neurons. Expression was correlated with methylation and accessibility across promoters and known enhancers. Several classes of transcription factor binding sites were enriched at regions of differential methylation and accessibility, including many that respond to synaptic activity. Finally, both regions of differential methylation and those of differential accessibility showed a surprising >10-fold enrichment of explained heritability associated with addictive behavior, as well as schizophrenia- and neuroticism-associated regions, suggesting that common psychiatric illness is mediated through brain region-specific epigenetic marks.

## Introduction

Epigenetic modifications, and DNA methylation in particular, are strongly implicated in carrying information for stable transcriptional patterns in the brain^1^. DNA methylation is altered in the brain tissues of patients with neuropsychiatric disease, including schizophrenia^2-4^, Alzheimer’s^5^, and major depressive disorder^6^. Brain-specific functions have anatomical regional correlates and much of disease-based brain research is focused on identifying structures that mediate normal function, e.g. the hippocampus in memory^7^, the prefrontal cortex in cognition^8^, and the nucleus accumbens in addictive behavior^9^. A key priority for human normal and disease-based research is identifying the functional genomic differences among brain regions, including gene expression, DNA methylation and chromatin.

Previous studies comparing normal brain regions, including from our own group, have found few DNA methylation^10-12^ or gene expression^13^ differences among non-cerebellar brain regions. Genome-wide analyses of these brain tissues fail to sufficiently account for cellular heterogeneity making it difficult to identify brain region-specific signatures that contribute differentially to neurological diseases, although computational methods have attempted to bridge this gap^14-16^. In spite of this, two recent case/control schizophrenia studies^3,4^ identified DNA methylation signatures that differed among brain regions between the diseased and normal brain, though combined this represents less than 200 DMRs. In contrast, DNA methylation differences between glia and neurons have been widely reported^16-21^, but comparisons between these cell types within a single brain region is generally uninformative with regard to neuronal dysfunction in many disease states.

Here, we addressed this knowledge gap by comparing post-mortem samples, both bulk tissues and fractionated neuronal and glial nuclei, from four brain regions (prefrontal cortex (BA9), anterior cingulate gyrus (BA24), hippocampus, and nucleus accumbens (NAcc)) at base pair resolution using whole genome bisulfite sequencing (WGBS). We found substantial differences in DNA methylation between the neurons of individual brain regions at thousands of locations across the genome, particularly between the nucleus accumbens and the other three tissues. The magnitude of these differences is as dramatic as we and others have previously reported between tumor and normal cells^22-24^, and is in marked contrast to methylation differences comparing bulk tissue across brain regions. We also examined both gene expression and chromatin accessibility (via ATAC-seq^25^) in an independent set of sorted neuronal and glial nuclei isolated from the nucleus accumbens and prefrontal cortex. In comparing these datasets, we found that differential methylation, gene expression, and chromatin accessibility intersected over genes highly relevant for region-specific neuronal function (i.e. dopamine receptor signaling in the nucleus accumbens). Regions of differential methylation and/or accessibility showed significant enrichment of explained heritability for neurological traits, particularly neuroticism and schizophrenia. Together, these data represent, to our knowledge, the largest and most comprehensive genome-wide epigenetic analysis of sorted nuclei from functionally diverse brain tissues. Our results have broad implications for future brain epigenetic studies, and suggest that comparisons of neurons within experimental frameworks (e.g. case/control studies, developmental trajectories, or inter-region comparisons) are more informative than comparisons of neurons to glia.

## Results

### Cell type heterogeneity within and between specimens obscures epigenetic differences between brain regions

To better understand the epigenetic regulation of normal brain function, we have mapped the DNA methylation landscape using whole-genome bisulfite sequencing of four human post-mortem brain regions: dorsolateral prefrontal cortex (BA9), anterior cingulate cortex (BA24), hippocampus (HC), and nucleus accumbens (NAcc). Functional abnormalities within each of these brain regions have been implicated in many neurological disorders including schizophrenia, Alzheimer’s, and major depressive disorder (MDD)^26-31^. We collected samples from the 4 brain regions from multiple donors (Supplementary Table 1) and generated 27 sequencing libraries at an average coverage of 8.4x per sample (Supplementary Table 2). We focus on epigenetic differences on the autosomes. Principal component analysis of the WGBS data binned in 1 kb intervals revealed no clear segregation between the different brain regions (Figure 1a), consistent with previous reports^10-12^.

**Figure 1.**
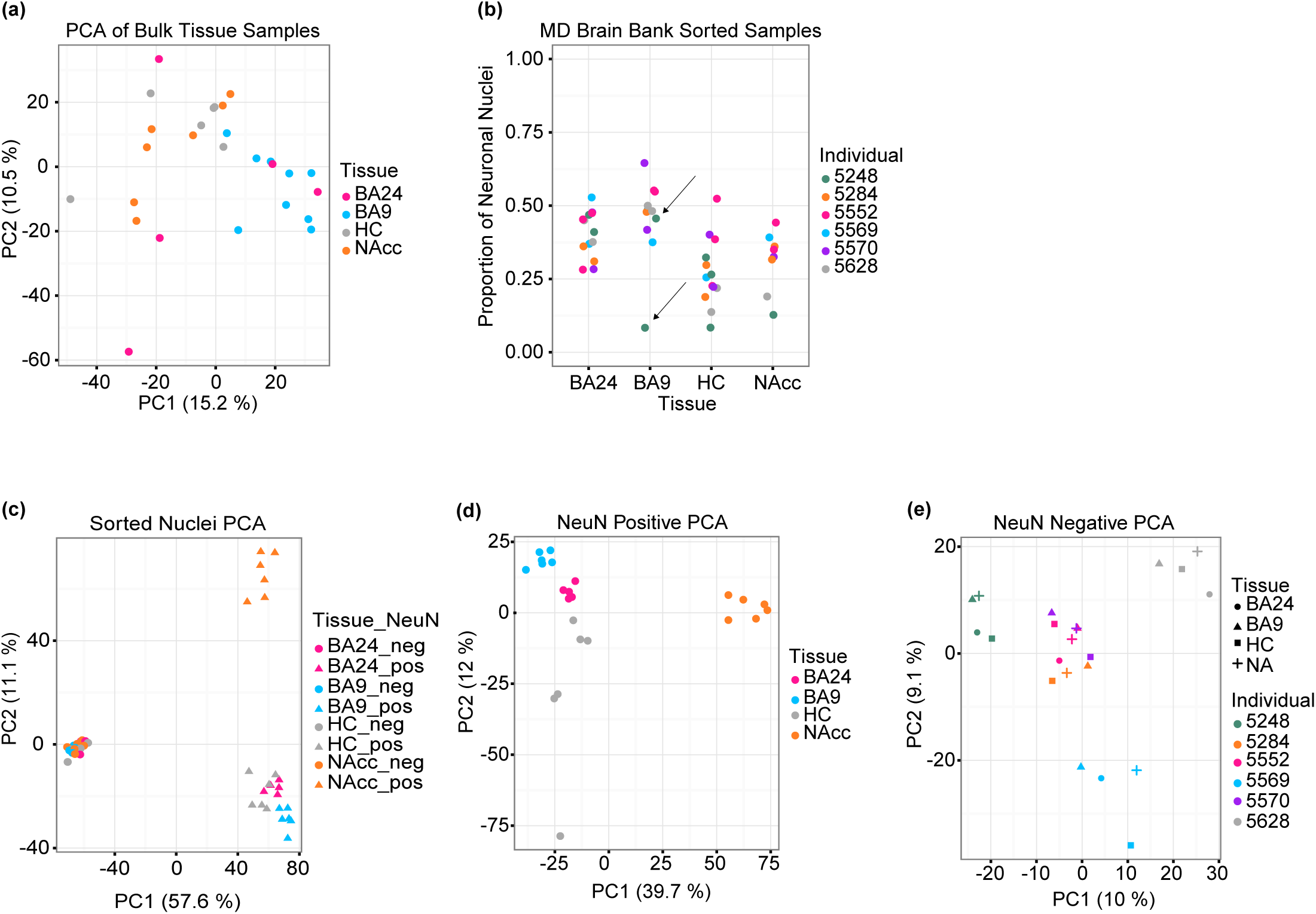
Differences in DNA methylation between brain regions are restricted to neuronal nuclei. (a) Principal component analysis of distances derived from average CpG methylation levels in 1 kb intervals along the autosomes of bulk brain tissue samples, assayed by whole-genome bisulfite sequencing (BA24 = anterior cingulate cortex, n = 5; BA9 = prefrontal cortex, n = 9; HC = hippocampus, n = 6; NAcc = nucleus accumbens, n = 7). Note the lack of segregation by tissue. (b) Sorting nuclei from bulk tissue samples revealed high variation in the proportion of neuronal nuclei isolated from distinct brain regions (BA24, n = 6; BA9, n = 6; HC, n = 6; NAcc, n = 6). Note that some pieces were split and sorted multiple times; arrows indicate an example of two punches from the same tissue sample. (c-e) Principal component analysis of distances derived from average CpG methylation levels binned in 1 kb intervals along the autosomes of (c) all sorted samples (BA24, n = 9; BA9, n = 12; HC, n = 12; NAcc, n = 12), (d) only neuronal nuclei (pos) (BA24, n = 5; BA9, n = 6; HC, n = 6; NAcc, n = 6) or (e) only glial nuclei (neg) (BA24, n = 4; BA9, n = 6; HC, n = 6; NAcc, n = 6) from each brain region.

We hypothesized that this lack of brain region segregation might be due to confounding by cell type heterogeneity. Indeed, changes in cell type composition have been reported in patients with major depressive disorder, schizophrenia, and bipolar disorder^32-35^ and across brain development^36^. To address this, we used fluorescence-activated nuclei sorting to separate neuronal nuclei from non-neuronal nuclei (hereafter referred to as glia) based on the nuclear neuronal marker NeuN (*RBFOX3*), followed by WGBS in a total of 45 samples from 6 donors at an average coverage of 11.1x (Supplementary Table 3; Supplementary Figure 1). Surprisingly, we observed that the proportion of isolated NeuN^+^ nuclei varies substantially within and between brain regions, as well as between different samplings from the same tissue specimen (Figure 1b). Indeed, principal component analysis of the sorted DNA methylation data now revealed clear segregation between brain regions and cell types (Figure 1c), showing that variation in cell type composition masks region-specific differences in DNA methylation. The first principal component represents cell type and explains 57% of the variation, while the second principal component represents a difference between the neurons of the NAcc and the neurons of the other three brain regions. In contrast to the differences between brain regions in the neuronal cell type, we observe no differences between brain regions in the glial cell type. To explore this further, we performed principal component analysis separately in the two cell types. This analysis showed distinct clustering of the 4 different brain regions in neurons (Figure 1d) whereas glial samples cluster by donor rather than brain region (Figure 1e). There are no clear brain region-specific patterns in autosomal global methylation, but we confirmed previous observations^19-21^ of higher global CpG and non-CpG methylation in neurons compared to glia (Supplementary Figure 2). These data clearly indicate that the neurons, rather than the glia, contain the relevant functional variability in DNA methylation between brain regions.

### Neuronal nuclei isolated from different brain regions display widespread differences in DNA methylation

To identify neuronal differentially methylated regions (DMRs), regions where at least one of the brain regions differed from at least one other region while accounting for the variation between biological replicates, we extended the approach of BSmooth^37^ to multi-group comparisons (Methods). We identified 13,074 autosomal neuronal DMRs containing 255,537 CpGs (1.1% of all CpGs analyzed), with a mean methylation difference of at least 10% between brain regions and family-wise error rate of 5% or less (by permutation). In contrast, we found only 114 autosomal DMRs using data from isolated glial nuclei and 71 autosomal DMRs using data from bulk tissue (Figure 2a; Supplementary Tables 4-6). This demonstrates that biologically relevant methylation differences between neurons from distinct brain regions are masked by the high proportion of glia across brain regions and the substantial variation of glial proportions across samples.

**Figure 2.**
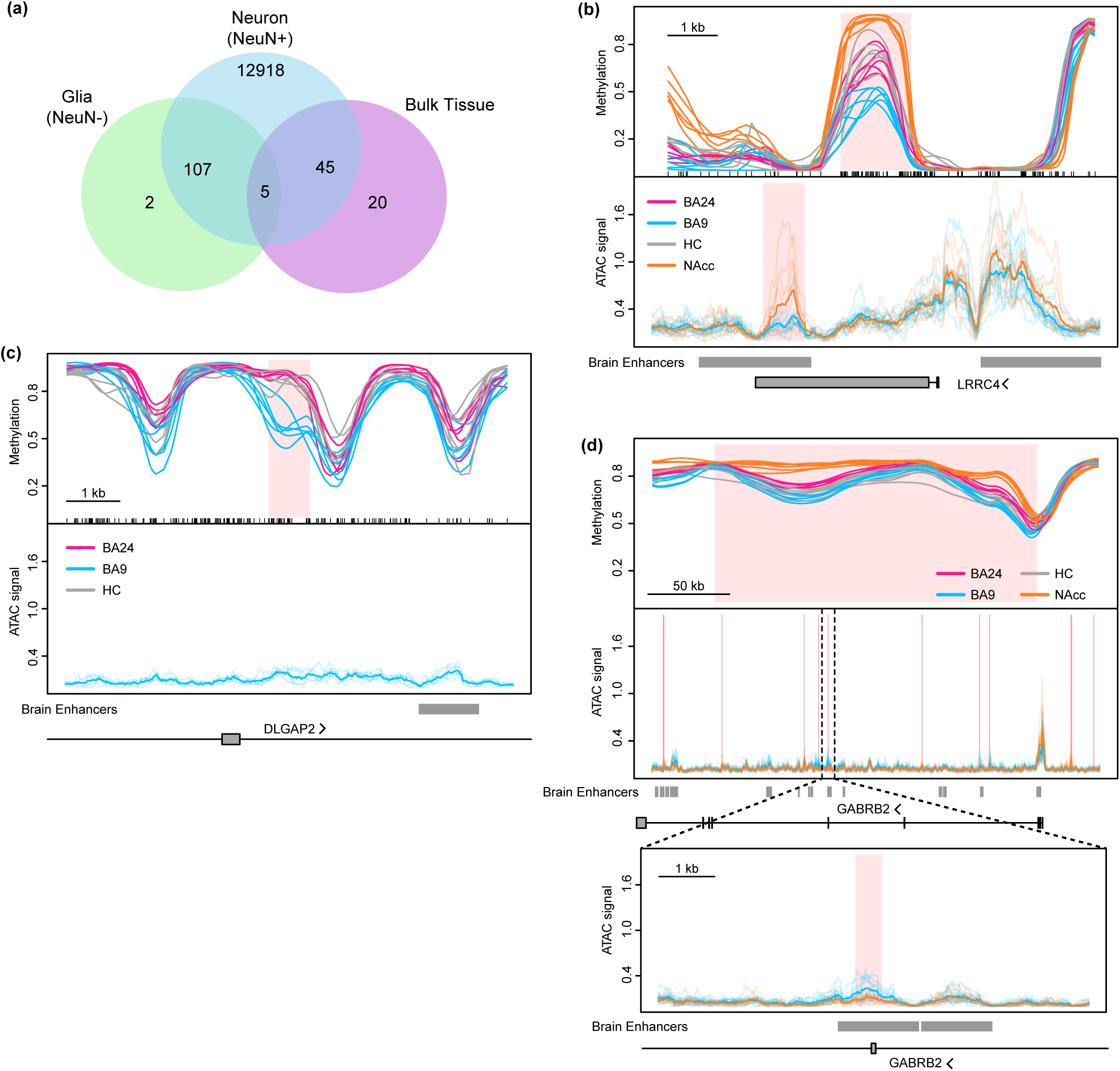
Neuronal nuclei isolated from different brain regions display widespread differences in DNA methylation. (a) Overlap of DMRs identified between brain regions within neurons, glia, and bulk tissues. (b-d) Methylation values for neuronal nuclei from the nucleus accumbens (orange, n = 6), hippocampus (grey, n = 6), anterior cingulate gyrus (BA24, pink, n = 5) and prefrontal cortex (BA9, blue, n = 6) from six individuals, as well as normalized ATAC sequencing coverage for neuronal nuclei from each of six individuals (transparent lines) in nucleus accumbens (n = 5) and prefrontal cortex (n = 6). Note: methylation and ATAC data are from different sets of individuals. Average ATAC coverage is indicated by opaque lines. Regions of differential methylation (DMRs) or differentially accessible ATAC peaks (DAPs) are shaded pink (see Methods). Overlap with brain-specific enhancers (see text) and protein-coding genes is depicted below each graph. Examples include: (b) a DMR between neurons in the 4 different brain regions (BA24, nWGBS = 6; BA9, nWGBS = 6, nATAC = 6; HC, nWGBS = 6; NAcc, nWGBS = 6, nATAC = 5); (c) a DMR between BA24, BA9 and hippocampus (BA24, nWGBS = 6; BA9, nWGBS = 6, nATAC = 6; HC, nWGBS = 6); (d) a large differentially methylated block across an entire gene containing multiple focal DAPs, one of which is expanded below (BA24, nWGBS = 6; BA9, nWGBS = 6, nATAC = 6; HC, nWGBS = 6; NAcc, nWGBS = 6, nATAC = 5).

To uncover the function of these DMRs, we computed overlaps and enrichment of these DMRs with common genomic features (Supplementary Figure 3). With respect to protein-coding genes, DMRs were most enriched in 3′ UTRs, with modest to no enrichment upstream of the transcription start site (TSS) or over the gene body. We found the expected enrichment of DMRs in CpG island shores and shelves^38,39^, over a large (215 Mb) set of enhancer-like regions mapped using H3K27ac in human brain regions^40^, as well in a focused set (12 Mb) of permissive enhancers identified across many cell types and tissues^41^. Finally, using a map of chromatin states in 4 brain regions similar to the ones we have profiled^42^, we found high enrichment in putative regulatory regions. Together, these data suggest that our brain region-associated DMRs are enriched in regulatory and enhancer-like elements.

Of the 13,074 neuronal DMRs, 11,895 involved the nucleus accumbens being different from all three of the other brain regions, consistent with our principal component analyses. Most (73%) of the NAcc neuronal DMRs were hypermethylated compared to the other three tissues. *LRRC4* (also known as *NGL-2*) is shown as an example of this (Figure 2b) and is known to play important roles in brain development, synapse formation, and differentiation of neurons and glia^43^. Analysis of hypermethylated DMRs using the Genomic Regions Enrichment of Annotations Tool (GREAT)^44^ showed enrichment in GO categories neurotrophin signaling and telencephalon development while hypomethylated DMRs showed enrichment in dopamine signaling and synaptic transmission (Supplementary Table 7). Focusing on DMRs overlapping promoters we used Enrichr^45^ to find that genes with hypermethylated promoters (1,284) were enriched for GO categories including positive regulation of GTPase activity, regulation of neuron differentiation, and synaptic transmission as well as KEGG pathways for glutamatergic and cholinergic synapse (Supplementary Table 8). Genes with hypomethylated promoters (264) were enriched in GO categories including adenylate cyclase-activating dopamine receptor signaling pathway and dopamine receptor signaling pathway and KEGG pathways for morphine addiction, dopaminergic synapse, and GABAergic synapse. These findings are consistent with the fact that 95% of NAcc neurons are GABAergic medium spiny neurons that express D1 or D2 dopamine receptors^46^ and the well documented role of the NAcc in addiction (reviewed in^9^).

Given the overwhelming differences between the NAcc and the other brain regions, we hypothesized that DMRs between the other three brain regions could be masked. Therefore, we repeated our analysis using only the neuronal samples from prefrontal cortex (BA9), anterior cingulate cortex (BA24), and hippocampus and identified 208 autosomal DMRs comprising 6,489 CpGs and 217,265 bp (Supplementary Tables 4, 9). One example where a DMR is present between these regions is at the *DLGAP2* locus (Figure 2c). DLGAP2 is a main component of postsynaptic scaffolding complexes important for proper synapse function, has been implicated in autism^47^ and Alzheimer’s^48^, and is differentially methylated in a rodent model of post-traumatic stress disorder ^49^. GREAT analysis of these DMRs showed enrichment for genes involved in nervous system development and neurogenesis among others (Supplementary Table 7). While we detected fewer methylation differences between these brain regions than when comparing them to the NAcc, these DMRs represent the largest set of differentially methylated loci found to date between normal, non-cerebellar brain tissues.

In addition to small DMRs (<11 kb), we also identified large blocks of differential methylation among neurons from these 4 brain regions. We discovered 1,808 differentially methylated blocks containing >1.7 million CpGs, with a median width of 64 kb, and mean methylation difference between brain regions greater than 10% (Supplementary Table 4; Supplementary Table 10). Several genes encoding GABA_A_ (α1, β2, β3, γ3) receptor subunits as well as the GABA_B2_ receptor were found in hypermethylated blocks in NAcc (example shown in Figure 2d). Interestingly, 23% of these blocks cover the entirety of a protein coding gene (411 blocks over 563 genes) (Supplementary Table 10); these genes showed enrichment for GO biological processes including neuron fate commitment, synaptic transmission, and regulation of neuron differentiation, as well as enrichment in KEGG pathways for enrichment in olfactory transduction, amphetamine addiction, and synaptic vesicle cycle (Supplementary Table 8).

### Neurons and glia have very divergent methylation profiles

In agreement with previous reports^16-21^, we found substantial methylation differences between neuronal and glial nuclei across all four brain regions. Simultaneously comparing all 45 samples, we identified 97,924 autosomal DMRs between cell types with mean methylation difference greater than 10% and large blocks of differential methylation (19,072 blocks with a median width of 48 kb) (Supplementary Tables 4, 11, 12). Several DMRs were found within *RBFOX3*, which encodes NeuN (the marker used to distinguish neuronal from non-neuronal nuclei) (Figure 3a). The entirety of *QKI*, a gene encoding an RNA-binding protein involved in oligodendrocyte differentiation^50^, is covered by a large block of differential methylation and is hypomethylated in glia (Figure 3b). The majority of both the small DMRs (63%) and differentially methylated blocks (81%) are hypermethylated in neurons, consistent with our and other’s results indicating slightly higher global methylation levels in neurons (Supplementary Figure 2). While previous studies have reported many differentially methylated loci between neurons and glia within the prefrontal cortex, hippocampus, and superior temporal gyrus^14,19,20^, we identified 21,802 novel DMRs (Supplementary Table 13). Interestingly, many of these novel regions correspond to genomic locations where the cell type difference is only present in one tissue, specifically the nucleus accumbens, indicating that inclusion of more diverse brain tissues can expand the current catalog of methylation differences between neurons and glia.

**Figure 3.**
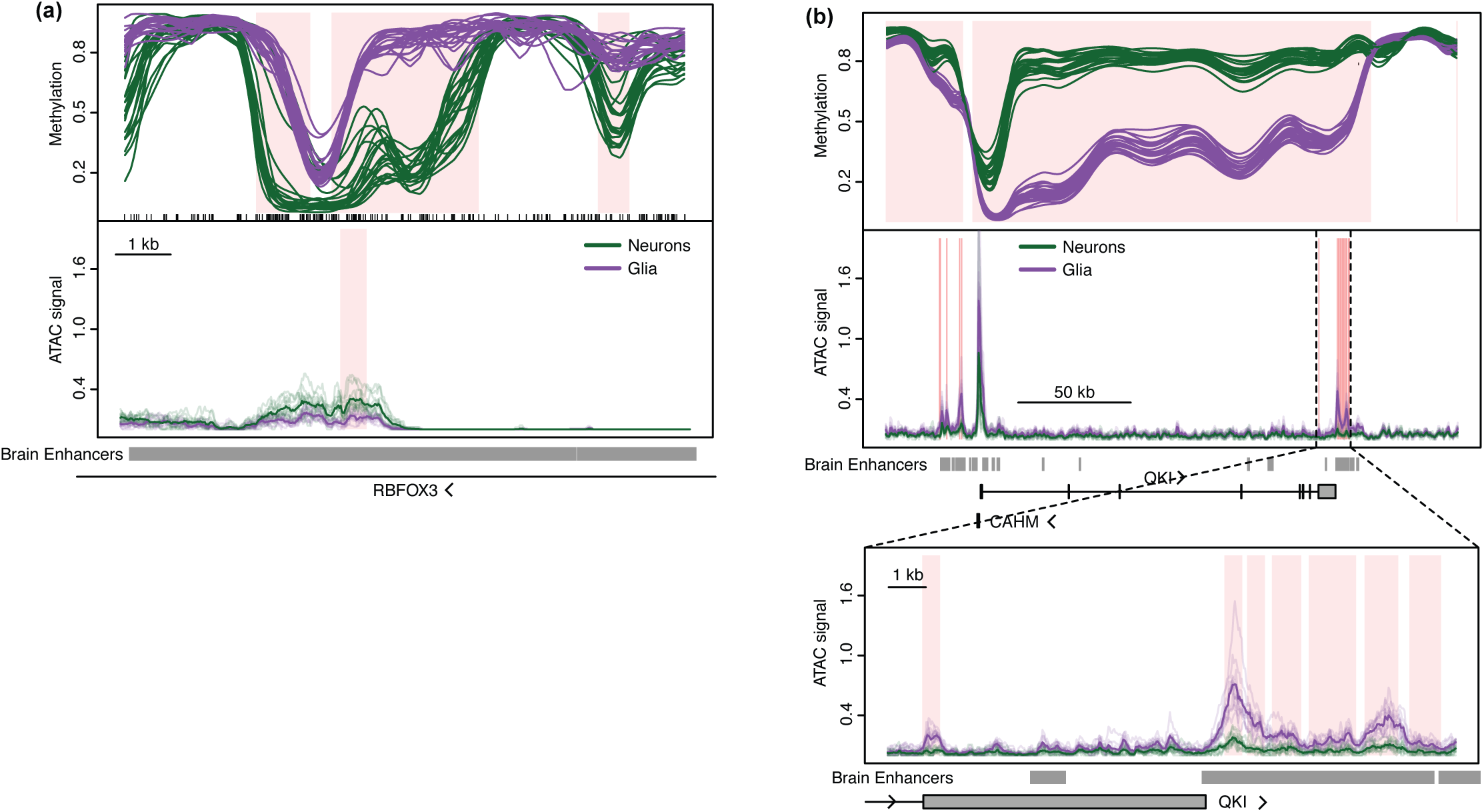
DNA methylation differs between neurons and glia.

Methylation values for neurons (green, n = 23) and glia (purple, n = 22) from four brain regions from six individuals. Normalized ATAC sequencing coverage for nuclei isolated from each of six individuals (transparent lines) in nucleus accumbens (n = 10) and prefrontal cortex (BA9, n = 12) is shown below each methylation plot. Average ATAC coverage is indicated by opaque lines. Regions of differential methylation (DMRs) or differentially accessible ATAC peaks (DAPs) are shaded pink (see Methods). Overlap with brain-specific enhancers (see text) and protein-coding genes is depicted below each graph. Examples include: (a) multiple small DMRs between neurons and glia and (b) a large differentially methylated block between neurons and glia containing multiple focal DAPs, one of which is expanded below.

### Gene expression and chromatin accessibility differ between neurons, but not glia, from the nucleus accumbens and prefrontal cortex

Given the enrichment of neuronal DMRs in transcriptional regulatory regions (e.g. enhancer-like regions and near TSSs) and the divergence between the NAcc and the other tissues, we sought to determine the relationship among these differentially methylated loci, gene expression, and DNA accessibility. We generated gene expression (RNA-seq) and chromatin accessibility (ATAC-seq^25,51^) data using samples from 6 additional donors (Supplementary Table 1), constituting an independent set of neuronal and glial nuclei isolated from the NAcc and prefrontal cortex (BA9), the two most divergent neuronal populations based on our methylation analyses.

We found 2,952 genes to be differentially expressed between NAcc and BA9 in neurons (FDR 5%). In contrast, there was 1 gene (*TENM3*) differentially expressed between NAcc and BA9 in glial cells supporting our conclusion of homogeneity in glial cells between brain regions (Figure 4a, b; Supplementary Tables 14, 15). Genes expressed at a higher level in BA9 (1,479) were enriched in GO biological processes including synaptic transmission, neuron projection guidance, and axon guidance. Given that the prefrontal cortex provides glutamatergic inputs to the NAcc^52^, it was unsurprising that the most enriched KEGG pathways were cAMP signaling pathway and glutamatergic synapse. In contrast, genes expressed at a higher level in the NAcc (1,473) were enriched in addiction-related GO biological processes including regulation of catecholamine secretion, adenylate cyclase-inhibiting G-protein coupled receptor signaling pathway, and response to amphetamine (Supplementary Table 8).

**Figure 4.**
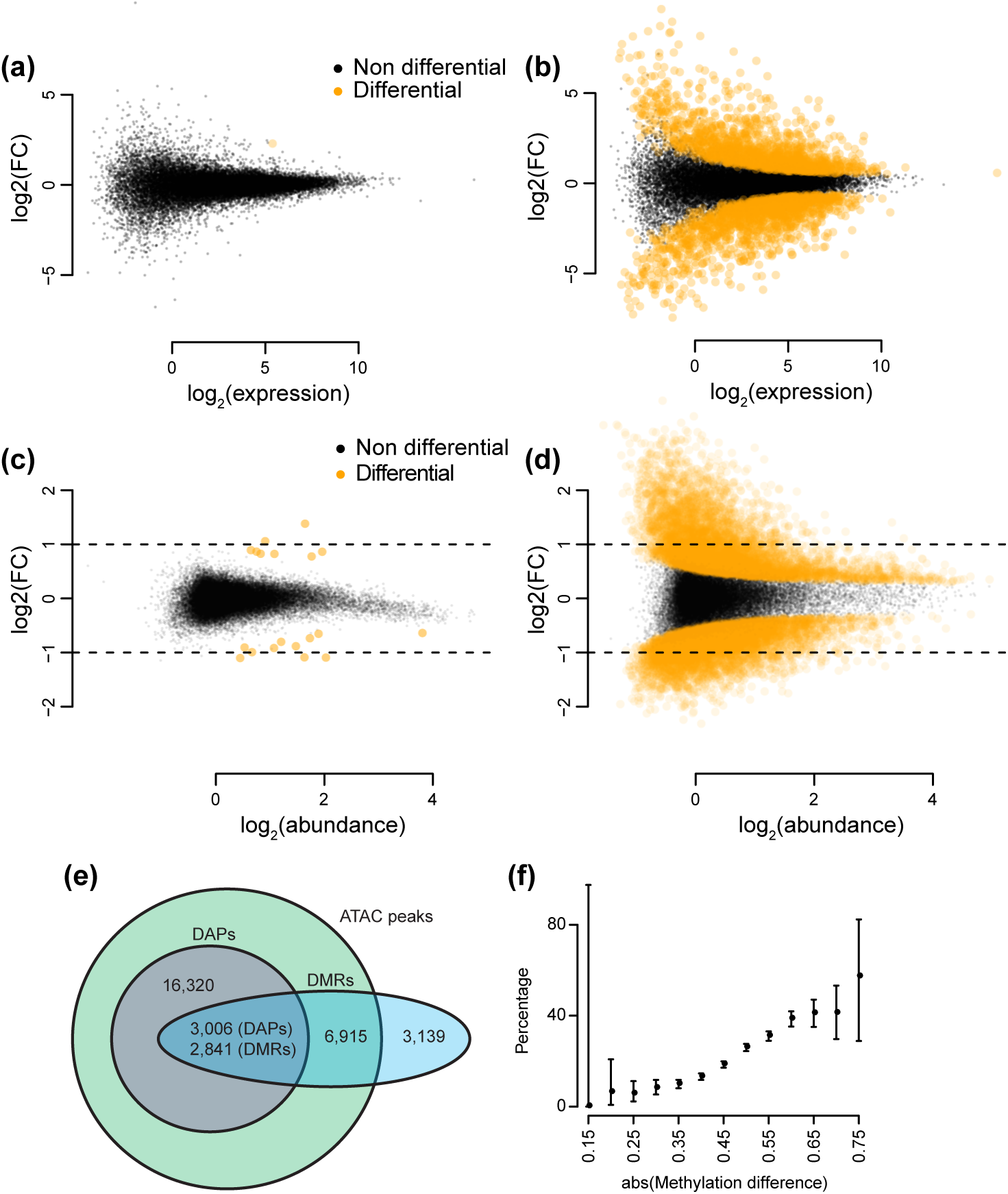
Chromatin accessibility and gene expression differ between brain regions in neurons, but not in glia. Mean-difference plots of gene expression data comparing nucleus accumbens (NAcc) to prefrontal cortex (BA9) in (a) glia (NAcc, n = 4; BA9, n = 5) and (b) neurons (NAcc, n = 5; BA9, n = 6). Differentially expressed genes (DEGs) are shown in orange (see Methods). Mean-difference plots of peak accessibility data comparing NAcc to BA9 in (c) glia (NAcc, n = 5; BA9, n = 6) and (d) neurons (NAcc, n = 5; BA9, n = 6). Differentially accessible peaks (DAPs) are those orange points that additionally have an absolute log fold change > 1 (see Methods). Data plotted in (c-d) were randomly down sampled (20% of DAPs, 10% of non-DAPs) to reduce over-plotting. (e) Overlap of neuronal DMRs between NAcc and BA9 with neuronal DAPs and ATAC peaks. (f) Estimate and 95% confidence interval for the percentage of neuronal DMRs that overlap a neuronal DAP (n = 2,841) as a function of the absolute difference in average methylation over the DMR (NAcc vs. BA9).

### Differential methylation and accessibility mark complementary parts of the genome

We measured chromatin accessibility via ATAC-seq and identified 424,597 peaks covering 324 Mb using peak calling on a pooled meta-sample (Methods). These ATAC peaks show the expected enrichment over 5’ UTRs, exons, promoters, FANTOM5 enhancers and other regulatory regions (Supplementary Figure 4a, b)

We performed a differential analysis of ATAC-seq peaks, accounting for biological variation, between NAcc and BA9 (Methods). For neurons we found 70,079 neuronal peaks (52 Mb) to be differentially accessible (FDR 5%), while for glia we found only 19 differentially accessible peaks (0.01 Mb) (Figure 4c, d; Supplementary Tables 16, 17; Methods). These results again strongly support our conclusion that glial cells are homogeneous between brain regions. Given that many differentially abundant peaks were associated with small fold changes between conditions, we focused on the 19,326 neuronal peaks covering 12 Mb that had an absolute log_2_(fold change) > 1 between NAcc and BA9, which we term differentially accessible peaks (DAPs).

These DAPs have a similar enrichment pattern as ATAC peaks when compared to random genomic regions (Supplementary Figure 4c). However, we reasoned that a better comparison would be to those ATAC peaks with little evidence of differential accessibility (null ATAC peaks, FDR > 5%). Surprisingly, this showed DAPs to be depleted in genic elements, enhancers, and other regulatory regions, with only slight enrichment in intergenic regions compared to null ATAC peaks (Supplementary Figure 4d-f). GREAT analysis of DAPs show enrichment in only one obvious brain-related GO category, glutathione derivative biosynthetic process; however, they are enriched in the MSigDB Pathway term for genes involved in GABA synthesis, release, reuptake and degradation, consistent with the predominance of GABAergic medium spiny neurons in the NAcc^46^ (Supplementary Table 8).

We next investigated the relationship between methylation and accessibility in our dataset. Of the 13,074 neuronal DMRs, 12,895 (11.7 Mb) involve differences between BA9 and NAcc and we focus on these DMRs for comparison with the neuronal DAPs. Most DMRs (76%) overlap an ATAC peak, but far fewer (22%) overlap a DAP (Figure 4e). Despite the fact that only 22% of DMRs overlap a DAP, they are strongly enriched in DAPs (log_2_(OR)=4.2, P < 2.2 × 10^−16^) and vice versa (log_2_(OR)=4.9, P < 2.2 × 10^−16^). Thus, differential methylation does not imply differential accessibility. However, we do find that greater methylation difference increases the chance of differential accessibility (Figure 4f). Further, if a region exhibits both differential methylation and accessibility, the two measures are highly concordant with 99.9% of these DMRs having higher methylation when the region is less accessible. If DMRs and DAPs do not overlap, they are far apart: the median distance from a DMR to a DAP is 54 kb (109 kb from a DAP to a DMR). GREAT analysis shows that DAPs are enriched for GO categories that are less brain region-specific as compared to DMRs (Supplementary Table 8). Together, these data indicate that DMRs and DAPs mark distinct regions of the genome.

### DMRs and DAPs are enriched for explained heritability of GWAS traits

We hypothesized that the epigenetic differences we found between brain regions might mark regions important in neurological diseases. To answer this, we estimated their contribution to the amount of heritability explained in genome-wide association studies (GWAS) using stratified LD score regression^53^. This method is a more principled approach to the question of genetic importance of various genomic regions than considering enrichment of leading GWAS SNPs in the same regions.

We separately evaluated our different sets of regions (DAPs, DMRs, blocks). We compare these regions to the signal observed in conserved regions - the regions Finucane et al.^53^ found to carry the strongest signal - as well as to FANTOM5 enhancers. As a negative control, we first examined enrichment for height, HDL, LDL, and triglyceride levels. As expected, the regions marked by differential methylation and accessibility do not show significant enrichment of explained heritability for these traits, unlike conserved regions (Figure 5a, Supplementary Figure 5c). We also observed that rheumatoid arthritis, a disease we have previously found to be associated with methylation in whole blood^54^, does not display significant enrichment over our regions of interest (Supplementary Figure 5d).

**Figure 5.**
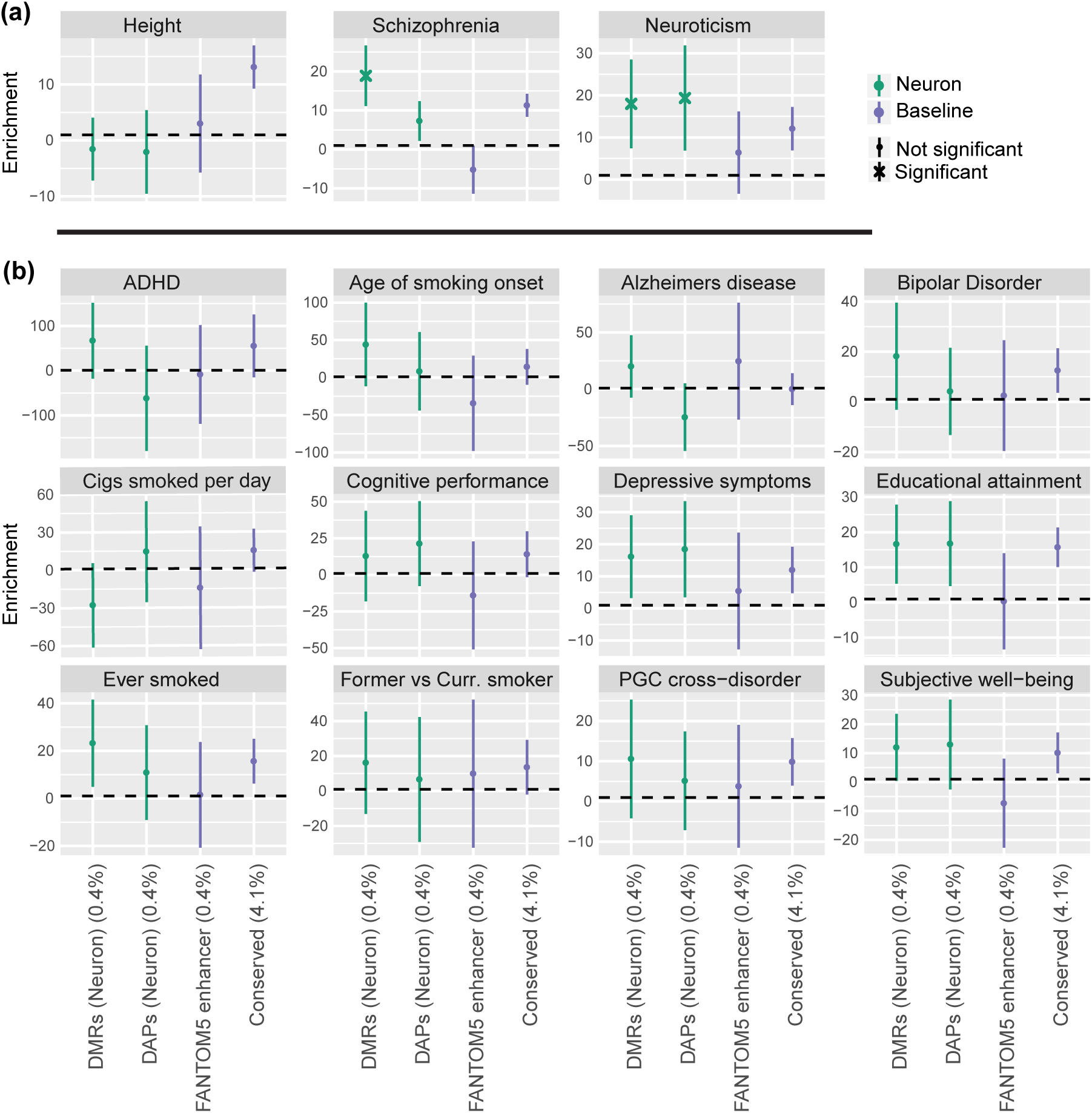
Neuronal DMRs and DAPs are highly enriched for explained heritability of neuropsychiatric GWAS traits. Estimates and 95% confidence intervals for the enrichment of explained heritability of GWAS traits (see Methods). Enrichments within neuronal DMRs and DAPs are contrasted with 1) enrichments within a set of permissive enhancers from the FANTOM5 project and 2) enrichments within regions of highly conserved sequence, which were previously shown to be highly enriched for explained heritability across a broad range of GWAS traits. The size of each category is reported as a percentage of the size of the autosomal genome. (a) Neuronal DMRs and DAPs are significantly enriched for explained heritability in neuroticism and schizophrenia GWAS traits (DMRs only); height is included as a negative control. (b) DMRs or DAPs in neuronal nuclei have > 10-fold enrichment for explained heritability in twelve neurological GWAS traits. In most cases, these enrichments are comparable to or larger than either baseline category, albeit it with wider confidence intervals due to the small genomic size of these features.

In marked contrast, examination of enrichment for neurological traits revealed striking results. We find significant enrichment (5% FDR) of one or more sets of regions for schizophrenia, neuroticism, educational attainment, PGC cross-disorder and ever smoked (Figure 5b, Supplementary Figure 5b). We find the largest enrichment values for DMRs and DAPs between brain regions within neurons, whereas DMRs and DAPs between neurons and glia are more often significant, but with a lower enrichment value. To interpret this, we note that (1) essentially all neuronal DMRs and DAPs between brain regions are also DMRs and DAPs between neurons and glia and (2) as a consequence of the LDSC method, the size of the query regions has a major impact on the uncertainty of explained heritability; neuronal DMRs and DAPs between brain regions are 2-6 times smaller than DMRs and DAPs between neurons and glia. Indeed, it is striking that neuronal DMRs and DAPs between brain regions have very large enrichment estimates across all 12 neurological traits (Supplementary Figure 5a).

De novo mutations have been hypothesized to contribute to the burden of some neurological traits. Recently, 10,387 de novo single nucleotide variants (SNVs) were identified in 192 autistic children with unaffected parents^55^. These mutations are slightly enriched in our neuronal DMRs (log2(OR)=0.80, P < 0.0001 by simulation, Supplementary Figure 5e), however, so too are de-novo mutations detected in a Dutch control population^56^ (log2(OR)=0.56, P < 0.0015 by simulation, Supplementary Figure 5f). This suggests that these neuronal DMRs may be hotspots for de novo mutations though not necessarily specific to autism.

### The relationship between differential expression, methylation and accessibility over promoters and enhancers

We next examined the relationship between differential expression, differential methylation and differential accessibility. To do so requires linking DMRs and DAPs to individual genes which is straightforward when they overlap genic promoters (+/− 2 kb of the transcription start site) or gene bodies. Selecting all genes with a DMR in their promoter, we find expression and methylation to be negatively correlated, as expected (Figure 6b), although the degree of correlation depends on the location of the DMR with respect to the transcriptional start site, with highest correlation 2 kb downstream of the TSS (Figure 6a). Extending to the entire gene body only slightly decreased the correlation (Figure 6c). Performing the same analysis with differential chromatin accessibility, we found that accessibility was positively correlated with differential expression, suggesting that the level of differential accessibility quantitatively influences gene expression (Figure 6a, d). Accessibility had peak correlation right at the TSS and substantially lower correlation within the gene body (Figure 6a, e). There is a slight interaction between accessibility and methylation in the sense that genes containing both a DAP and a DMR in their promoter show slightly stronger correlation between these marks and gene expression (Supplementary Figure 6).

**Figure 6.**
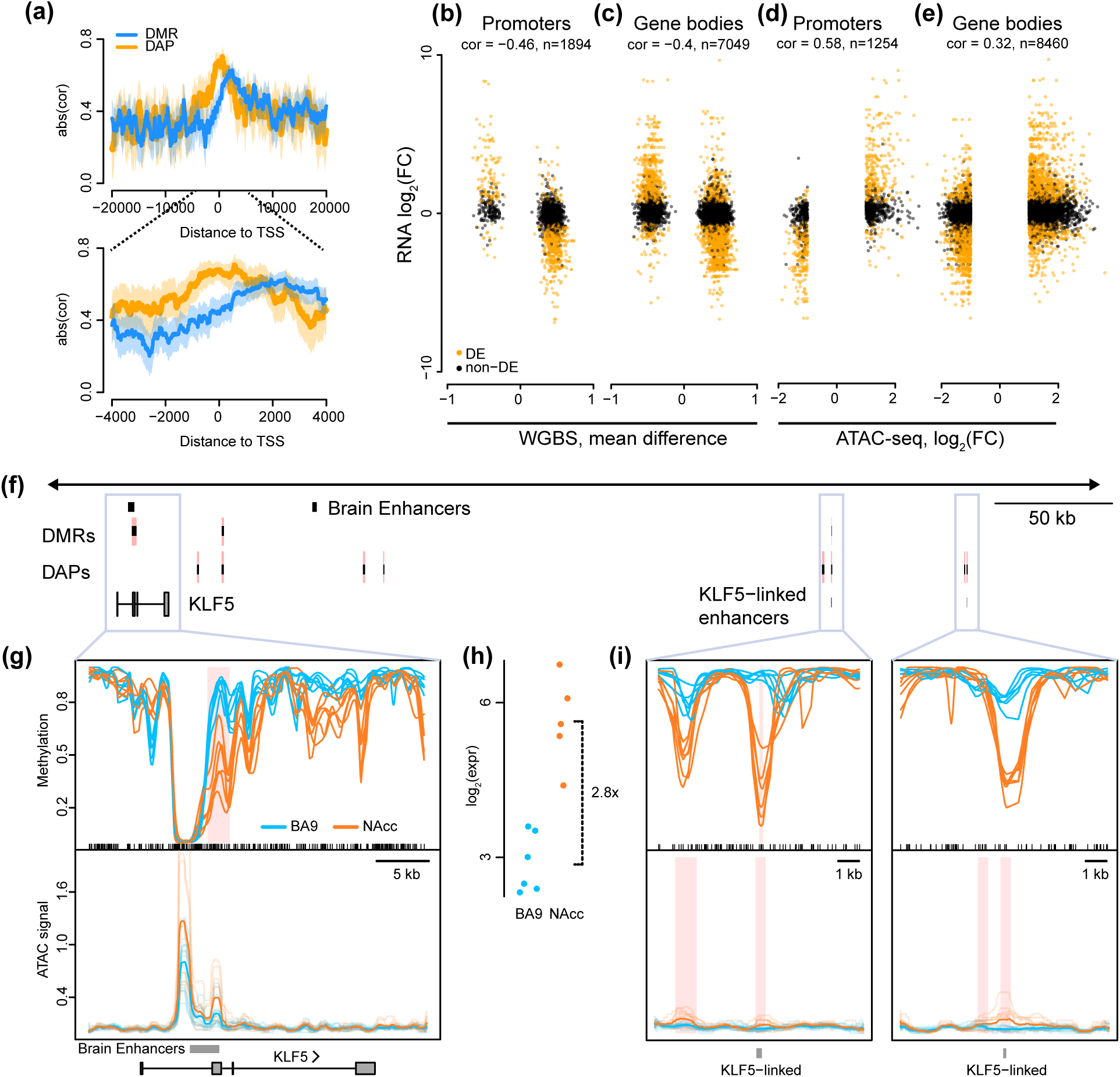
The relationship between differential expression, methylation, and accessibility in neurons over promoters and enhancers. (a) Absolute Pearson correlation between expression and methylation (blue) or accessibility (yellow) centered around the transcriptional start site in comparison of neurons from nucleus accumbens (NAcc) and prefrontal cortex (BA9). Estimated correlation is indicated by opaque lines and 95% confidence interval indicated by shaded region. (b)-(e) Scatterplots showing Pearson correlation of differential expression with differential methylation (b,c) or accessibility (d,e) in promoters (b,d) and gene bodies (c,e) in comparison of neurons from nucleus accumbens (NAcc) and prefrontal cortex (BA9). Note that the methylation measurements were obtained from samples distinct from accessibility and expression measurements. Differentially expressed genes are shown in orange. (f) Genome view of a 400 kb region containing KLF5 illustrating the relationship between gene expression, DMRs, and DAPs around protein-coding genes and linked enhancers (see text and Methods). The directionality of all DMRs and DAPs linked to KLF5 are consistent with the direction of differential expression (see text). (g,i) Methylation values and normalized accessibility for neuronal nuclei from the nucleus accumbens (NAcc, n_WGBS_ = 6, n_ATAC_ = 5; orange) and prefrontal cortex (BA9, n_WGBS_ = 6, n_ATAC_ = 6; blue) for six individuals, around (g) KLF5 and (i) two distant enhancers. (h) Gene expression of KLF5 in nucleus accumbens and prefrontal cortex, along with estimated log fold change of expression between the two brain regions in neurons (NAcc, n_RNA_ = 5, orange; BA9, n_RNA_ = 6, blue).

To expand our analysis to include enhancers, we used data from the FANTOM5 project, which linked 11,607 genes to 27,451 enhancer regions using CAGE data across more than five hundred samples^41^. While not brain-specific, using their data we can link 8,361 putative enhancer regions to 1,686 differentially expressed genes between NAcc and BA9. Just over half (4,237) of these putative enhancer regions overlap an ATAC peak, and 59% of these putative enhancers overlap a gene body (including introns). We next investigated the joint effect of differential methylation and accessibility in promoters and enhancers on these 1,686 enhancer-linked, differentially expressed genes. Of these, 945 do not have a DMR or a DAP overlapping either a promoter or a linked enhancer. There are 373 enhancer-linked differentially expressed genes where only their promoter overlaps a DMR or DAP; the epigenetic state of 86% of these genes is consistent with gene expression, in that genes with higher expression are more accessible and less methylated. The remaining genes have at least one DAP or DMR in a linked enhancer. Many (66%) of these genes are fully consistent with a simple model: all DMRs and DAPs overlapping either enhancers or promoters of the gene are more accessible and less methylated in the same brain region, and this brain region shows higher expression. An example of this is the KLF5 gene which has one DMR and two DAPs in two linked enhancers (Figure 6f-i).

### Differential methylation and accessibility mark genes involved in brain region-specific activity

Many neuronal DMRs and DAPs occur near or within differentially expressed genes involved in axon guidance. One such gene is *SATB2* which is hypermethylated in the NAcc with decreased expression compared to BA9 (Figure 7a). SATB2 is a transcriptional repressor that targets many genes involved in axon guidance ^57,58^. Two SATB2 targets, *BCL11B* and DCC, contain hypomethylated DMRs and are more highly expressed in NAcc consistent with the repressive function of SATB2. (Supplementary Tables 5, 14). Another example is the semaphorin family member SEMA7A. Semaphorins play important roles throughout development in establishing neuronal connectivity networks (reviewed in ^59^). We found *SEMA7A* to be hypermethylated in NAcc with decreased expression compared to BA9 (Figure 7b).

**Figure 7.**
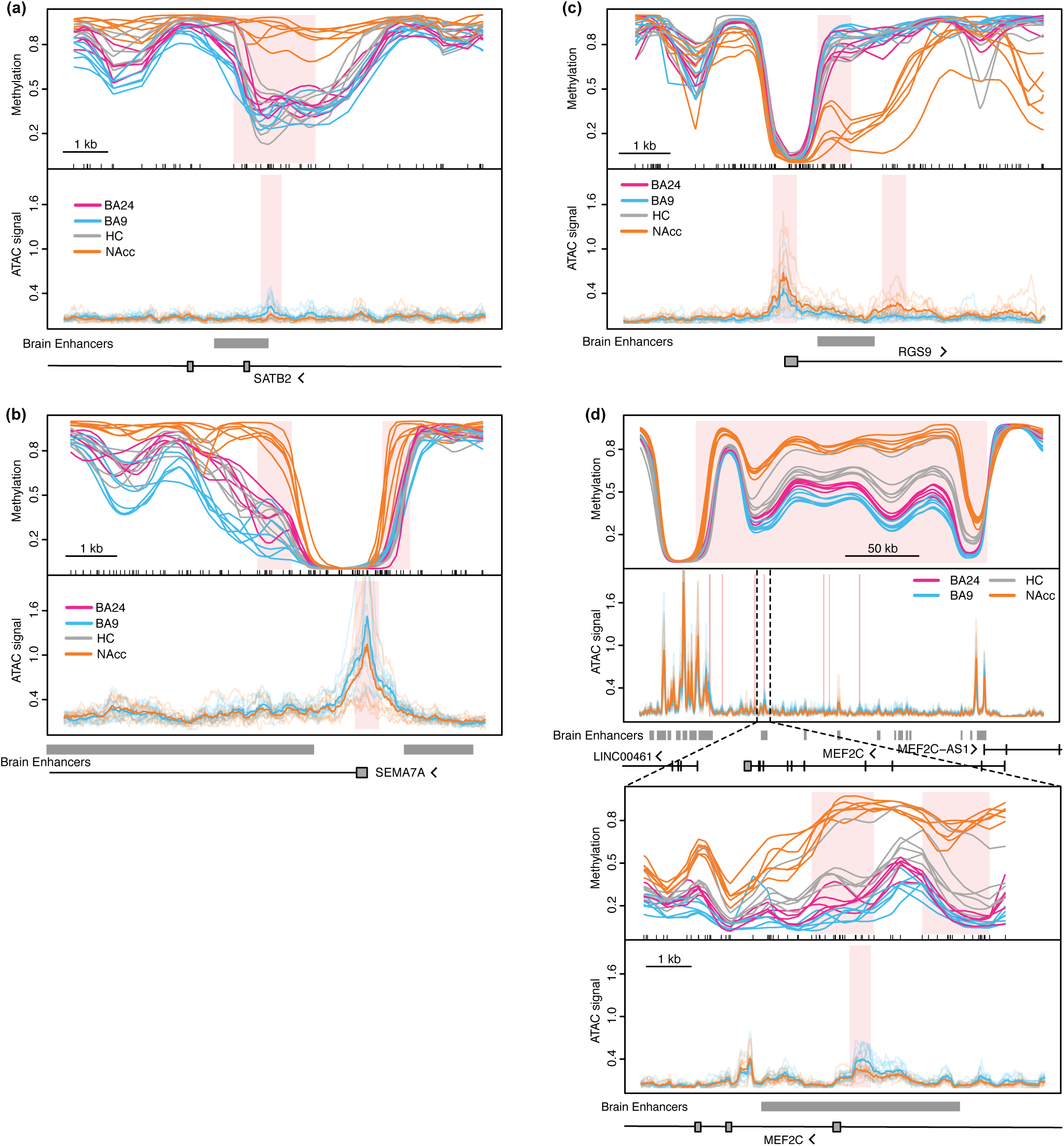
Methylation differences between neuronal nuclei from different brain regions overlap enhancers and genes involved in brain region-specific functions. Methylation values for neuronal nuclei from the nucleus accumbens (orange), hippocampus (grey), anterior cingulate gyrus (BA24 (pink) and prefrontal cortex (BA9) (blue) for six individuals, as well as normalized ATAC sequencing coverage for neuronal nuclei from each of 6 individuals (transparent lines) in nucleus accumbens and prefrontal cortex (BA24, nWGBS = 6; BA9, nWGBS = 6, nATAC = 6; HC, nWGBS = 6; NAcc, nWGBS = 6, nATAC = 5). Note: methylation and ATAC data are from different sets of individuals. Average ATAC coverage is indicated by opaque lines. Regions of differential methylation (DMRs) or differentially accessible ATAC peaks (DAPs) are shaded pink (see Methods). Overlap with brain-specific enhancers (see text) and protein-coding genes is depicted below each graph. Examples include: (a) a small DMR in the SATB2 gene overlapping both an enhancer and a DAP; (b) a DAP in SEMA7A flanked by two small DMRs and two enhancers; (c) a small DMR in RGS9 overlapping an enhancer and flanked by DAPs; (d) a large differentially methylated block overlapping MEF2C containing multiple focal DAPs and enhancers, one of which overlaps a small DMR and is expanded below.

In addition, many members of the regulator of G protein signaling (RGS) family of proteins are differentially methylated and differentially expressed in NAcc (*RGS6, RGS8, RGS9, RGS14*, and *RGS20*); an example is given in Figure 7c, *RGS9*. These proteins are primarily expressed in the brain and regulate neurotransmitter release, synaptic plasticity, and synaptic transmission (reviewed in ^60^). Expression of *RGS9* is 60-fold higher in NAcc, is important in addiction (cocaine, morphine), and has been implicated in schizophrenia and dyskinesias^60^. Another family member, RGS6, which is hypermethylated and downregulated in NAcc, is known to interact with DNMT1 complex member, DMAP1, to inhibit the repressive activity of this complex^61^. Both *SEMA7A* and *RGS9* illustrate the aforementioned relationship whereby a change in accessibility over a promoter is accompanied by downstream changes in DNA methylation, both of which are consistent with changes in gene expression (Figure 7b, c).

Another gene, *MEF2C*, encodes a synaptic activity-regulated transcription factor that restricts the number of synapses and dendritic spines in medium spiny neurons of the NAcc and has been identified as a possible autism spectrum disorder risk gene^62-65^. This entire gene is highly methylated in NAcc containing multiple small DMRs and is included in a large hypermethylated block (Figure 7d). In addition, *MEF2C* is downregulated 11-fold in NAcc compared to BA9. These are just a few examples of the many genes with brain region-specific functions whose epigenome differs between NAcc and BA9.

### Differential epigenetic regulation of transcription factors and their binding sites

Given the importance of the epigenome in regulating transcription factor (TF) binding, we used Haystack^66^ to identify motifs enriched in DMRs and DAPs. Our DMRs were enriched in CpG-containing motifs whose methylation influences TF binding^67-70^ (Figure 8a), and many TFs whose motifs were enriched were also differentially expressed between NAcc and BA9 neurons (Supplementary Figure 7a). DMRs that overlap promoters (Figure 8b; Supplementary Table 20) and those located elsewhere in the genome (Supplementary Figure 7b; Supplementary Table 21) were enriched for binding sites of synaptic activity-regulated TFs. These include multiple MEF2 family members (MEF2A, MEF2C, and MEF2D) and immediate-early genes (IEGs) (*FOS, JUN, EGR1-3*). Using MEF2C ChIP-seq ENCODE data from a lymphoblastoid cell line, we found that 181 neuronal DMRs (143 hypermethylated and 38 hypomethylated in NAcc) and 626 differentially expressed genes overlap a MEF2C binding site. These sites include known MEF2C targets *JUN*^71^ and *BDNF* which are both downregulated in NAcc, consistent with the hypermethylation and downregulation of *MEF2C* itself. IEGs with enriched motifs include EGR2 and EGR3, implicated in schizophrenia^72^ and bipolar disorder^73^, respectively, and AP-1 complex components (c-Jun, Fos, and JDP family members) with known roles in addiction^74-78^. A recent study performing ATAC-seq on mouse dentate granule neurons found that AP-1 binding sites were enriched in regions of chromatin that became open after neuronal activation^79^. We also observe enrichment within our DAPs, regardless of the direction of accessibility (*i.e*. more accessible in NAcc or more accessible in BA9) suggesting differential sites of TF binding between brain regions (Figure 8c; Supplementary Figure 7c).

**Figure 8.**
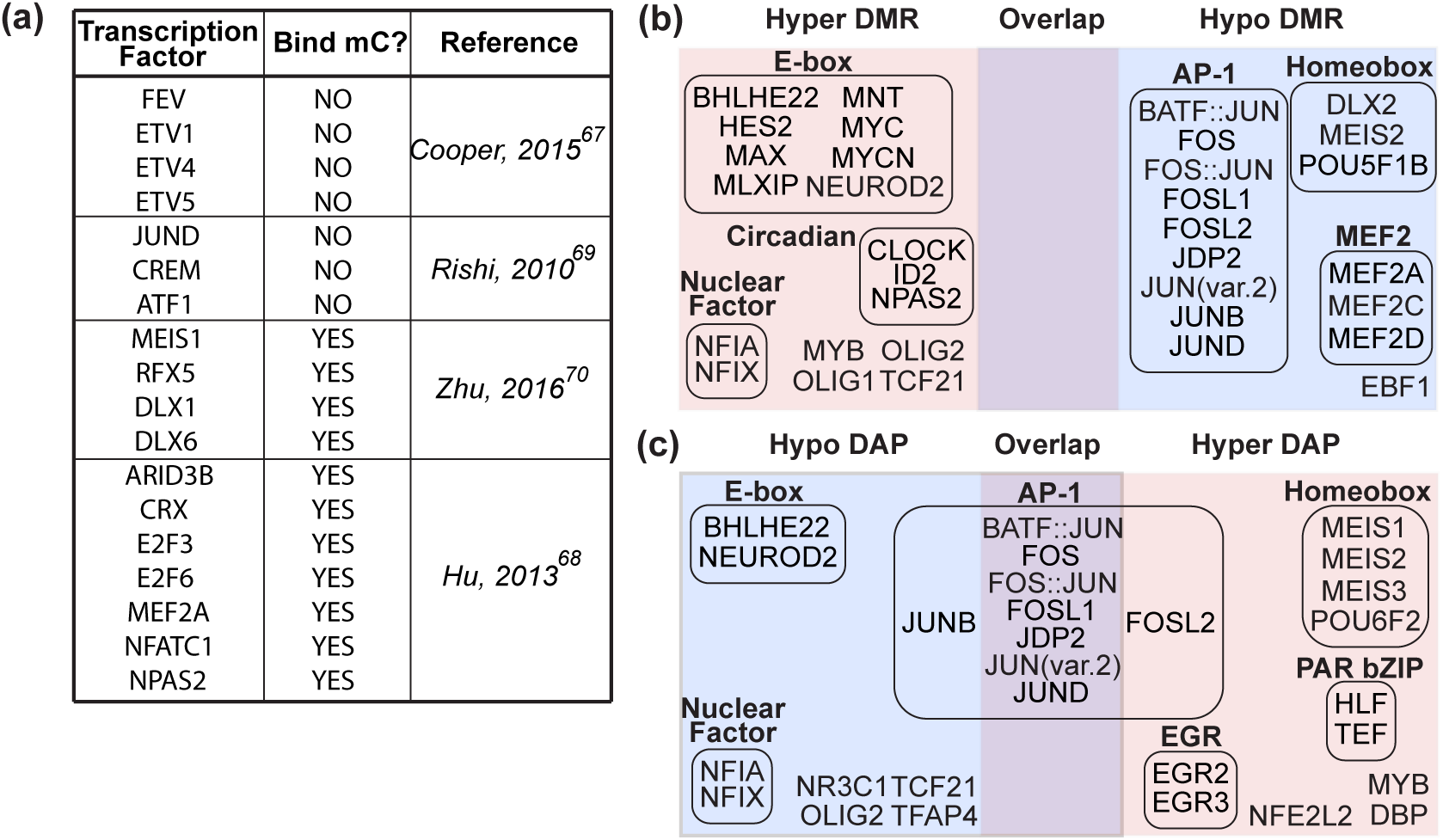
Transcription factor motifs enriched in DMRs and DAPs that overlap promoters. (a) Transcription factors with motifs enriched in our DMRs and/or DAPs (regardless of overlap with promoters) whose binding is impacted by CpG methylation (see Methods). (b,c) Transcription factor motifs enriched in promoter-specific (b) DMRs and (c) DAPs. Only motifs corresponding to transcription factors expressed in BA9 and NAcc neuronal nuclei are shown. “Hyper DMR” and “Hypo DMR” refer to regions of differential methylation when comparing NAcc to BA9. Similarly, “Hyper DAP” and “Hypo DAP” refer to ATAC peaks that are more, or less accessible, respectively, in NAcc vs. BA9. Transcription factors are roughly grouped by family (or function).

Other DMRs were enriched in binding sites for TFs important during brain development many of which were also enriched in “hypo DAPs” (*i.e*. regions more accessible in BA9 than NAcc) (Figure 8b, c). For example, NEUROD2 activates genes required for neurogenesis, migration, and axon guidance of cortical projection neurons^80^, and its reduced expression and hypermethylation of its recognition sites in NAcc is consistent with its important role in the cortex (BA9). Taken together, our data show that differential methylation and accessibility mark regions of genome being regulated by TFs that are responsive to synaptic activity and suggest that these TFs likely regulate different targets in each tissue.

## Discussion

Our results reveal unexpectedly high variation in neuron/glia ratios across the brain, even in the same small specimen. This variation greatly confounds the search for brain region-specific molecular patterns. We find that DNA methylation, as well as gene expression and chromatin accessibility, are remarkably consistent in glia across brain regions, while neuron-specific differences are as substantial as we find in other tissues, contrary to many published papers, including our own^10-12^. As previously reported^14,16-21^, we find substantial differences between neurons and glia. Most of the differentially methylated regions we found between neurons from distinct brain regions distinguish nucleus accumbens from the other brain regions. Given that GABAergic medium spiny neurons comprise 95% of nucleus accumbens neurons and that the other brain regions display more neuronal heterogeneity, we hypothesize that further cell fractionation will enhance detection of DMRs in other brain regions. This is supported by a recent methylation profiling of three neocortical neuron subtypes in mice, which revealed more than twice as many differentially methylated regions compared to a mixed neuronal sample^81^. Our neuronal DMRs were most enriched in enhancer regions and 3’ UTRs emphasizing the importance of DNA methylation in regulatory regions.

Surprisingly, we found that regions of differential methylation and differential accessibility were largely non-overlapping, indicating that differential methylation does not imply differential accessibility. This shows the power of profiling both methylation and accessibility. Gene expression was negatively correlated with methylation and positively correlated with chromatin accessibility as expected. Interestingly, this correlation was strongest over the TSS for DAPs while for DMRs it was strongest ∼2-4 kb into the gene body. Accessibility and methylation over known enhancers are consistent with each other and with gene expression, even between different enhancers for the same gene.

Additionally, both neuronal DMRs and DAPs display >10-fold enrichment of explained heritability associated with addictive behavior (e.g. ever smoked), consistent with the well-characterized role of the nucleus accumbens in addiction^9^, as well as an enrichment of DMRs for schizophrenia- and neuroticism-associated regions. Surprisingly, the DMRs are between brain regions, yet they are enriched for multiple psychiatric GWAS traits, in support of the conclusion that disease mechanisms are brain region-specific. We find that regions of hypomethylation in the nucleus accumbens were enriched in genes involved in dopaminergic pathways including *DRD1*, which encodes D1 dopamine receptor, and addiction mechanisms (e.g. *MEF2C* and *RGS9*) suggesting that the epigenome promotes specialized neuronal function.

Finally, we found very strong enrichment for several classes of transcription factor binding sites at DMRs and DAPs, including many (*MEF2C, FOS, JUN, EGR1-3*) that respond specifically to synaptic activity (reviewed in^82^) and several which are reported to be methylation sensitive. These data are consistent with a link between synaptic activity and long-term epigenetic modification of transcription factor binding sites. This finding agrees with previous data in adult mouse dentate gyrus neurons, suggesting that DNA methylation could perpetuate transient stimuli into long-lasting neuronal plasticity^83^. Together, these results are consistent with the notion that insofar as epigenetics is involved in human disease, it may be related to disrupted developmental and phenotypic plasticity^84^.

## Acknowledgements

This work was supported by funding awarded to A.P.F. (U01MH104393n) through the enhanced Genotype-Tissue Expression (eGTEx) project supported by the Common Fund of the Office of the Director of NIH. We would like to thank Hao Zhang from the Flow Cytometry Cell Sorting Core Facility at Johns Hopkins School of Public Health for flow sorting. The core facility is supported by CFAR: 5P30AI094189-04, 1S10OD016315-01, and 1S10RR13777001.

## Contributions

L.F.R, K.D.H, and A.P.F designed the study; L.F.R. performed nuclei sorting, DNA and RNA extractions; V.R.D. performed ATAC-seq; R.T., A.I., C.M.C. performed WGBS and RNA-seq library preparation and sequencing; A.P.F. oversaw the experiments; K.D.H. oversaw the data analysis. L.F.R, P.F.H., K.D.H, and A.P.F. performed data analysis and interpreted the results; L.F.R, P.F.H., K.D.H, and A.P.F wrote manuscript.

## Corresponding authors

Correspondence to: Andrew P. Feinberg or Kasper D. Hansen

## ONLINE METHODS

### Data availability

Raw and processed data generated during this study are available on the Gene Expression Omnibus (GSE96615).

## Experimental Methods

### Human Postmortem Brain Samples

Fluorescence-activated nuclei sorting was performed on flash-frozen postmortem dorsolateral prefrontal cortex (BA9), hippocampus (HC), nucleus accumbens (NAcc), and anterior cingulate gyrus (BA24) from six individuals not affected with neurological or psychiatric disease. These samples underwent nuclei extraction and sorting as described below for subsequent DNA methylation analysis. Additionally, neuronal nuclei were isolated from the nucleus accumbens (NAcc) and dorsolateral prefrontal cortex (BA9) of 6 different individuals for RNA-seq and ATAC-seq analysis. To underscore the importance of cell sorting, we also prepared DNA from unsorted material from the four brain regions above (BA9, n=9; HC, n=7; NAcc, n=7; BA24, n=5). The majority of individuals were matched between sorted and unsorted, but not all. All samples were obtained from the University of Maryland Brain and Tissue Bank which is a Brain and Tissue Repository of the NIH NeuroBioBank (Supplementary Table 1). This study was approved under IRB00061004.

### Nuclei Extraction, Cell Sorting, and DNA Isolation

Total nuclei were extracted via sucrose gradient centrifugation as previously described^1^ with the following changes. For WGBS analysis, a total of 2 × 250 mg of frozen tissue per sample was homogenized in 5 mL of lysis buffer (0.32 M sucrose, 10 mM Tris pH 5 mM CaCl_2_, 3 mM Mg acetate, 1 mM DTT, 0.1 mM EDTA, 0.1% Triton X-100) by douncing 50 times in a 40 mL dounce homogenizer. Lysates were combined and transferred to a 38 mL ultracentrifugation tube and 18 mL of sucrose solution (1.8 M sucrose, 10 mM Tris pH 8.0, 3 mM Mg acetate, 1 mM DTT) was dispensed to the bottom of the tube. The samples were then centrifuged at 28,600 rpm for 2 h at 4°C (Beckman Optima XE-90; SW32 Ti rotor). After centrifugation, the supernatant was removed by aspiration and the nuclear pellet was resuspended in 500 uL staining mix (2% normal goat serum, 0.1% BSA, 1:500 anti-NeuN conjugated to AlexaFluor488 (Millipore, cat#: MAB377X) in PBS) and incubated on ice. Unstained nuclei and nuclei stained with only secondary antibody served as negative controls. The fluorescent nuclei were run through a Beckman Coulter MoFlo Cell Sorter with proper gate settings (Supplementary Figure 1). A small portion of the NeuN^+^ and NeuN^−^ nuclei were re-run on the sorter to validate the purity which was greater than 95%. Immuno-negative (NeuN^−^) and -positive (NeuN^+^) nuclei were collected in parallel. Sorted nuclei were pelleted by adding 2 mL of sucrose solution, 50 uL of 1 M CaCl_2_, and 30 uL of Mg acetate to 10 mL of nuclei in PBS. This solution was incubated on ice for 15 min, then centrifuged at 3,000 rpm for 20 min. The nuclear pellets were flash frozen in liquid nitrogen and stored at −80°C. DNA was extracted from the frozen nuclear pellets using the MasterPure DNA Extraction kit (Epicentre, Madison, Wisconsin, USA) following the manufacturer’s instructions.

### Whole genome bisulfite sequencing (WGBS)

WGBS single indexed libraries were generated using NEBNext Ultra DNA library Prep kit for Illumina (New England BioLabs, Ipswich, MA, USA) according to the manufacturer’s instructions with modifications. 400 ng gDNA was quantified by Qubit dsDNA BR assay (Invitrogen, Carlsbad, CA, USA) and 1% unmethylated lambda DNA (cat#: D1521, Promega, Madison, WI, USA) was spiked in to measure bisulfite conversion efficiency. Samples were fragmented to an average insert size of 350 bp using a Covaris S2 sonicator. Size selection was performed using AMPure XP beads and insert sizes of 300-400 bp were isolated (0.4x and 0.2x ratios). Samples were bisulfite converted after size selection using EZ DNA Methylation-Gold Kit (cat#: D5005, Zymo, Irvine, CA, USA) following the manufacturer’s instructions. Amplification was performed after the bisulfite conversion using Kapa Hifi Uracil+ (cat#: KK282, Kapa Biosystems, Boston, USA) polymerase using the following cycling conditions: 98°C 45s / 8cycles: 98°C 15s, 65°C 30s, 72°C 30s / 72°C 1 min. Final libraries were run on 2100 Bioanalyzer (Agilent, Santa Clara, CA, USA) High-Sensitivity DNA assay; samples were also run on Bioanalyzer after shearing and size selection for quality control purposes. Libraries were quantified by qPCR using the Library Quantification Kit for Illumina sequencing platforms (cat#: KK4824, KAPA Biosystems, Boston, USA), using 7900HT Real Time PCR System (Applied Biosystems). Libraries were sequenced with the Illumina HiSeq2500 using 125 bp paired-end single indexed run and 10% PhiX spike-in.

### Assay for transposase-accessible chromatin using sequencing (ATAC-seq)

NeuN^+^ and NeuN^−^ nuclei were isolated as previously described and 100,000 nuclei were used for ATAC-seq library preparation as per standard protocols^2^. Briefly, nuclei were suspended in lysis buffer and incubated 20 min on ice followed by centrifugation for 10 min as previously described. The transposition reaction was incubated for 1 h at 37°C (Nextera DNA library prep kit; cat #:FC-121-1031, Illumina). After PCR amplification of libraries and column clean up via the Qiagen MinElute PCR purification kit (cat#:28004, Qiagen, Valencia, CA, USA), an additional clean up with AMPure XP beads (0.8x ratio) was performed twice with 80% ethanol washes before quantification using a DNA High Sensitivity chip on a 2100 BioAnalyzer (Agilent, Santa Clara, CA, USA). Libraries were sequenced with the Illumina HiSeq4000 using 70 bp paired-end single indexed run with a 5% PhiX spike-in.

### RNA sequencing (RNA-seq)

RNA isolated from bulk tissue was assessed and only tissues with a RIN ≥ 4 were used for nuclei isolation. NeuN^+^ and NeuN^−^ nuclei were isolated as previously described with the addition of 20 U/mL RNAse Inhibitors (cat#: N8080119, Applied Biosystems) to the lysis buffer, sucrose solution, and antibody solution while protease inhibitor cocktail (cat#: 50-751-7359, Amresco) was added to the lysis buffer only. Approximately 200,000 nuclei were sorted directly into RLT buffer + 150 mM 2-mercaptoethanol and RNA was isolated using the Qiagen RNeasy Kit (cat #:74106, Qiagen, Valencia, CA, USA). Nuclear RNA quality was assessed by running samples on a Total RNA Pico Chip on a 2100 BioAnalyzer (Agilent, Santa Clara, CA, USA). RNA-seq libraries were created using 2.5 ng input RNA with the SMARTer^®^ Stranded Total RNA-Seq Kit - Pico Input Mammalian (cat#: 635005, Takara Bio, Mountain View, CA, USA) following the manufacturer’s instructions for degraded RNA samples. Libraries were sequenced with the Illumina HiSeq4000 using 70 bp paired-end single indexed run with 5% PhiX spike-in.

## Computational Methods

### Annotation

The hg19 build of the human reference genome was used for all analyses. Only analyses of autosomal data are reported. Genes, exons, introns, and UTRs were generated from GENCODE v19 (http://www.gencodegenes.org/releases/19.html)^3^ and CpG islands were downloaded from UCSC (http://genome.ucsc.edu/)^4,5^. CpG shores are defined as 2 kb flanking CpG islands and CpG shelves are defined as 2 kb flanking CpG islands. Promoters were defined as 4 kb centered on the transcription start site. The 15-state ChromHMM model for 7 adult brain tissues from the Roadmap Epigenomics Project^6^ was downloaded using the R/Bioconductor AnnotationHub package (v2.6.4).

### Whole genome bisulfite sequencing (WGBS)

#### Mapping and quality control of WGBS reads

We trimmed reads of their adapter sequences using Trim Galore! (v0.4.0) (http://www.bioinformatics.babraham.ac.uk/projects/trim_galore/) and quality-trimmed using the following parameters: `trim_galore -q 25 ‐‐paired ${READ1} ${READ2}`. We then aligned these trimmed reads to the hg19 build of the human genome (including autosomes, sex chromosomes, mitochondrial sequence, and lambda phage (accession NC_001416.1) but excluding non-chromosomal sequences) using Bismark^7^ (v0.14.3) with the following alignment parameters: `bismark ‐‐bowtie2 -X 1000 −1 ${READ1} −2 ${READ2}`. Supplementary Tables 2 and 3 summarize the alignment results. Using the reads aligned to the lambda phage genome, we estimated that all libraries had a bisulfite conversion rate > 99%.

We then used `bismark_methylation_extractor` to summarize the number of reads supporting a methylated cytosine and the number of reads supported a unmethylated cytosine for every cytosine in the reference genome. Specifically, we first computed and visually inspected the M-bias^8^ of our libraries. Based on these results, we decided to ignore the first 5 bp of read1 and the first 10 bp of read2 in the subsequent call to `bismark_methylation_extractor with parameters: ‐‐ignore 5 ‐‐ ignore_r2 10`. The final cytosine report file summarizes the methylation evidence at each cytosine in the reference genome.

#### Smoothing WGBS

We used BSmooth to estimate CpG methylation levels as previously described^8^. Specifically, we ran a ‘small’ smooth to identify small DMRs (smoothing over windows of at least 1 kb containing at least 20 CpGs) and a ‘large’ smooth to identify large-scale blocks (smoothing over windows of at least 20 kb containing at least 500 CpGs). Following smoothing, we analyzed all CpGs that had a sequencing coverage of at least 1 in all samples (n = 45 for sorted data, n = 27 for unsorted data).

#### Identification of small DMRs and large-scale blocks

Previously, we have used BSmooth to perform pairwise (two-group) comparisons^9^. In the present study, we had up to 8 groups to compare: 4 brain regions (BA9, BA24, HC, NAcc) and, for the sorted data, 2 cell types (NeuN^+^, NeuN^−^). Rather than running all 28 pairwise comparisons, we extended the BSmooth method to handle multi-group comparisons, which we refer to as the F-statistic method.

For the F-statistic method, we constructed a design matrix with a term for each group (e.g., BA9_neg for NeuN^−^ cells from BA9, BA9_pos for NeuN^+^ cells from BA9, etc.). For each CpG, we then fitted a linear model of the smoothed methylation levels against the design matrix. To improve standard error estimates, we thresholded the residual standard deviations at the 75% percentile and smoothed these using a running mean over windows containing 101 CpGs. We then combined the estimated coefficients from the linear model, their estimated correlations, and the smoothed residual standard deviations to form F-statistics to summarize the evidence that methylation differs between the groups at each of the CpGs.

Next, we identified runs of CpGs where the F-statistic exceeded a cutoff and where each CpG was within a maximum distance of the next. Specifically, we used cutoffs of F = 4.6^2^ for DMRs and F = 2^2^ for blocks (following^10^) and required that the CpGs were within 300 bp of one another for DMRs and 1000 bp of one another for blocks. For blocks, we also required that the average methylation in the block varied by at least 0.1 across the groups. These runs of CpGs formed our candidate DMRs and blocks. Each candidate DMR and block was summarized by the area under the curve formed when treating the F-statistic as a function along the genome (`areaStat`). We used permutation testing to assign a measure of statistical significance to each candidate DMR/block. We randomly permuted the design matrix, effectively permuting the sample labels, and repeated the F-statistic analysis with the same cutoffs using the permuted design matrix, resulting in a set of null DMRs/blocks for each permutation. We performed 1000 such permutations. We then asked, for each candidate DMR/block, in how many permutations did we see a null DMR/block anywhere in the genome with the same or better `areaStat` as the candidate DMR/block; dividing this number by the total number of permutations gives a permutation P-value for each DMR/block. Since we are comparing each candidate block/DMR against anything found anywhere in the genome in the permutation set, we are also correcting for multiple testing by controlling the family-wise error rate. Those candidates DMRs/blocks with a permutation P-value ≤ 0.05 form our set of DMRs/blocks.

#### Annotation of small DMRs and blocks

The F-statistic approach allows us to jointly use all samples for the identification of DMRs and blocks. However, it does not tell us which group(s) are hypomethylated or hypermethylated for the region. To assign such labels to our F-statistic DMRs and blocks, we used a post-hoc analysis for specific pairwise comparisons of interest: NeuN^+^ vs. NeuN^−^; NeuN^+^ cells in NAcc vs. NeuN^+^ cells in BA9, BA24, and HC; NeuN^+^ cells in NAcc vs. NeuN^+^ cells in BA9. We identified small DMRs and blocks using the original t-statistic method of BSmooth; an F-statistic DMR was assigned a label (e.g., hypermethylated in NeuN+ and hypomethylated in NeuN^−^) if the corresponding t-statistic DMR or block overlapped at least 50% of the F-statistic DMR or block. This procedure does not change the coordinates of the DMR/block and means an F-statistic DMR/block may be assigned multiple labels.

#### Subset analyses

We found, as expected, that the differences between NeuN^+^ and NeuN^−^ samples dominated our results (98,420 / 100,875 F-statistic DMRs and 19,072 / 20,373 F-statistic blocks were assigned the label, NeuN^+^ vs NeuN^−^; Supplementary Tables 11 and 12). To better focus on the differences between brain regions *within* a given cell type (NeuN^+^ or NeuN^−^), we repeated the F-statistic analysis using just the NeuN^+^ or NeuN^−^samples (Supplementary Tables 5, 6, 10). We again found that one group dominated: 11,895 / 13,074 F-statistic NeuN^+^ DMRs were specific to NAcc. To better focus on the differences between the remaining brain regions, we repeated the analysis using just the BA9, BA24, and HC NeuN+ samples (Supplementary Table 9).

#### Novel NeuN^+^ vs NeuN^−^ DMRs

Three published datasets of NeuN^+^ vs NeuN^−^ methylation differences^1,11,12^ were used for comparison with our DMRs. Data from Montano et al. was generated from our own lab and is accessible through GEO series accession number GSE48610. Differentially methylated sites from Kozlenkov et al. were obtained by request directly from the authors. DMRs from Lister et al. were obtained from http://brainome.ucsd.edu/BrainMethylomeData/CG_DMR_lists.tar.gz and converted to hg19 using the UCSC liftOver tool^13^. As three different platforms were used to measure methylation (CHARM, WGBS, and 450K, respectively), we combined the differentially methylated CpGs from the autosomes of each study and compared to the differentially methylated CpGs within our NeuN^+^ vs NeuN^−^ DMRs. Sites that were unique to our NeuN^+^ vs NeuN^−^ DMRs were reported as novel.

### Assay for transposase-accessible chromatin using sequencing (ATAC-seq)

#### Mapping and quality control of ATAC-seq reads

We trimmed reads of their adapter sequences using trimadap (v0.1, https://github.com/lh3/trimadap/archive/0.1.zip) with the following parameters: `trimadap-mt -3 CTGTCTCTTATACACATCTCCGAGCCCACGAGA ${READ1}; trimadap-mt -3 CTGTCTCTTATACACATCTGACGCTGCCGACGA ${READ2}`. We then aligned these trimmed reads to the hg19 build of the human genome (including autosomes, sex chromosomes, mitochondrial sequence, unplaced sequence, and unlocalized sequence) using Bowtie2^14^ (v2.2.5) with alignment parameters: `bowtie2 - X 2000 ‐‐local ‐‐dovetail`. Potential PCR duplicate reads were marked using MarkDuplicates from the Picard library (http://broadinstitute.github.io/picard/; v2.2.1). Supplementary Table 18 summarizes the alignment results for the 22 libraries.

#### Identifying differentially accessibly ATAC-seq peaks (DAPs)

Peaks were called using MACS^15^ (v2.1.0) on a metasample formed by combining all non-duplicate-marked reads with a mapping quality > 30 from the 22 samples: macs2 callpeaks ‐‐nomodel ‐‐nolambda ‐‐call-summits -t ${BAMS[@]}. We excluded those peaks overlapping the ENCODE mappability consensus blacklist regions (http://hgdownload.cse.ucsc.edu/goldenPath/hg19/encodeDCC/wgEncodeMapability/) and the blacklist for ATAC-seq created by Buenrostro et al.^2^ (https://sites.google.com/site/atacseqpublic/atac-seq-analysis-methods/mitochondrialblacklists-1). We extended +/- 250 bp from the summit of peak and merged overlapping peaks to form our initial set of peaks comprising 961,916 autosomal peaks.

For each sample, we counted the number of *fragments* (fragment = start of read1 to end of read2) overlapping each of the 961,916 peaks using the summarizeOverlaps() function in the GenomicAlignments R/Bioconductor package^16^ (v1.10.0). Specifically, we only counted those fragments where both reads had a mapping-quality score > 30, reads not marked as potential PCR duplicates, and those where any part of the fragment overlapped exactly one peak.

We then analyzed these data using the voom method, originally designed for differential expression analysis of RNA-seq data^17^. Briefly, the read counts were transformed to counts per million (cpm) and only those 424,597 / 961,916 peaks with at least 1 cpm for at least 5 samples (the size of the smallest group of samples) were retained. These 424,597 peaks were used in all downstream analyses described in the main text. We normalized these counts using TMM^18^, then used edgeR^19^ (v3.16.5) and limma^20^ (v3.30.7) to transform these counts to log_2_-cpm, estimate the mean-variance relationship, and compute appropriate observation-level weights ready for linear modelling.

In our design matrix, we blocked on donor (donor1, …, donor6) and included a term for each group (e.g., BA9_neg for NeuN^−^ cells from BA9, BA9_pos for NeuN^+^ cells from BA9, etc.). We ran surrogate variable analysis^21^ using the sva (v3.22.0) R/Bioconductor package and identified 4 surrogate variables, one of which correlated with the date on which these samples were flow-sorted. We ultimately decided to include all 4 surrogate variables in the linear model. Using the empirical Bayes shrinkage method implemented in limma, we tested for differential accessibility of peaks in three comparisons: (1) NAcc vs. BA9 in NeuN^+^ cells; (2) NAcc vs. BA9 in NeuN^−^ cells; (3) NeuN^+^ cells vs NeuN^−^ cells. For an ATAC-seq peak to be called a differentially accessible peak (DAP), it had to have a Benjamini-Hochberg adjusted P-value < 0.05 and an absolute log2 fold change > 1, the latter filter to focus on the regions that were plausibly more interesting and interpretable.

### RNA sequencing (RNA-seq)

#### Mapping and quality control of RNA-seq reads

We trimmed the first 3 bp of read1, which were derived from template switching oligos and not the cDNA of interest, using seqtk (https://github.com/lh3/seqtk; v1.2-r94) with the following parameters: `seqtk trimfq -b 3 ${READ1}`. We then quasi-mapped these trimmed reads to a FASTA file of protein-coding and lncRNA genes from GENCODE v19 (http://www.gencodegenes.org/releases/19.html)^3^ and performed transcript-level quantification using Salmon^22^ (v0.7.2). Supplementary Table 19 summarizes these results for the 20 libraries.

#### Identifying differentially expressed genes (DEGs)

We used tximport^23^ (v1.2.0) to compute normalized gene-level counts from the transcript-level abundance estimates (scaling these using the average transcript length over samples and the library size). Only autosomal genes with at least 1 cpm in at least 4 libraries (the size of the smallest group of samples) were retained for downstream analysis (24,161 / 33,351 genes). We normalized these counts using TMM^18^ then used edgeR^19^ (v3.16.5) and limma^20^ (v3.30.7) to transform these counts to log_2_-cpm, estimate the mean-variance relationship, and compute appropriate observation-level weights ready for linear modelling.

In our design matrix, we blocked on donor (donor1, …, donor6) and included a term for each group (e.g., BA9_neg for NeuN^−^ cells from BA9, BA9_pos for NeuN^+^ cells from BA9, etc.). We ran surrogate variable analysis^21^ using the sva (v3.22.0) R/Bioconductor package and identified 5 surrogate variables, some of which correlated with the date on which these samples were flow-sorted. We ultimately decided to include all 5 surrogate variables in the linear model. Using the empirical Bayes shrinkage method implemented in limma, we tested for differential expression of genes in three comparisons: (1) NAcc vs. BA9 in NeuN^+^ cells; (2) NAcc vs. BA9 in NeuN^−^ cells; (3) NeuN^+^ cells vs NeuN^−^ cells. For a gene to be called a differentially expressed gene (DEG), it had to have a Benjamini-Hochberg adjusted P-value < 0.05 with no minimum log2 fold change cutoff.

### Enrichment of DMRs, ATAC peaks, and DAPs in genomic features

#### Enrichment odds ratios and P-values

We formed a 2×2 contingency table of (n_11_, n12, n_2_1, n_22_); specific values of (n_11_, n_12_, n_21_, n_22_) are described below. The enrichment log odds ratio was estimated by log_2_(OR) = log2(n_11_) + log2(n_22_) − log2(n_12_) − log2(n_21_), its standard error was estimated by se(log2(OR)) = sqrt(1 / n_11_ + 1 /n_12_ + 1 / n_21_ + 1 / n_22_), and an approximate 95% confidence interval formed by [log_2_(OR) − 2 × se(log2(OR)), log_2_(OR) + 2 × se(log2(OR))]. We also report the P-value obtained from performing Fisher’s exact test for testing the null of independence of rows and columns in the 2×2 table (i.e. the null of no enrichment or depletion) using the `fisher.test()` function from the ‘stats’ package in R^24^.

### DMRs

For DMRs, we computed the enrichment of CpGs within DMRs inside each genomic feature (e.g., exons, FANTOM5 enhancers, etc.). Specifically, for each genomic feature, we constructed the 2×2 table (n_11_, n_12_, n_21_, n_22_), where:

- n_11_ = Number of CpGs in DMRs that were inside the feature
- n_12_ = Number of CpGs in DMRs that were outside the feature
- n_21_ = Number of CpGs not in DMRs that were inside the feature
- n_22_ = Number of CpGs not in DMRs that were outside the feature

The total number of CpGs, n = n_11_ + n_12_ + n_21_ + n_22_, was the number of autosomal CpGs in the reference genome covered by at least one read. We counted CpGs rather than DMRs or bases because this accounts for the non-uniform distribution of CpGs along the genome and avoids double-counting DMRs that are both inside and outside the feature.

### ATAC peaks

For ATAC peaks, we computed the enrichment of bases within ATAC peaks inside each genomic feature. Specifically, for each genomic feature, we constructed the 2×2 table (n_11_, n_12_, n_21_, n_22_), where:

- n_11_ = Number of bases in ATAC peaks that were inside the feature
- n_12_ = Number of bases in ATAC peaks that were outside the feature
- n_21_ = Number of bases in the rest of the genome that were inside the feature
- n_22_ = Number of bases in the rest of the genome that were outside the feature

The total number of bases, n = n_11_ + n_12_ + n_21_ + n_22_, was the number of autosomal bases in the reference genome. We counted bases rather than number of ATAC peaks to account for the slight variation in ATAC peak width, the large variation in width for the ‘rest of the genome’ features, and to avoid double-counting ATAC peaks that were both inside and outside the feature.

### DAPs

For DAPs, we computed the enrichment of bases within DAPs inside each genomic feature in two ways. Firstly, for each genomic feature, we constructed the 2 × 2 table (n_11_, n_12_, n_21_, n_22_), where:

- n_11_ = Number of bases in DAPs that were inside the feature
- n12 = Number of bases in DAPs that were outside the feature
- n_21_ = Number of bases in the rest of the genome that were inside the feature
- n_22_ = Number of bases in the rest of the genome that were outside the feature

Secondly, for each genomic feature, we constructed the 2 × 2 table (n_11_, n_12_, n_21_, n_22_), where:

- n_11_ = Number of bases in DAPs that were inside the feature
- n_12_ = Number of bases in DAPs that were outside the feature
- n_21_ = Number of bases in null-peaks that were inside the feature
- n_22_ = Number of bases in null-peaks that were outside the feature

‘Null-peaks’ were those ATAC-seq peaks that were not differentially accessible between the relevant condition (NAcc and BA9 in NeuN^+^ cells) based on the peak having a Benjamini-Hochberg adjusted P-value > 0.05 in the analysis of differential accessibility. By comparing to null-peaks rather than the rest of the genome, we account for the nonuniform distribution of ATAC peaks along the genome.

We counted the number of bases rather than the number of ATAC peaks to account for the slight variation in peak width and to avoid double-counting ATAC peaks that were both inside and outside the feature.

### Linkage disequilibrium score regression (LDSC)

We used stratified linkage disequilibrium score regression (LDSC^25^) to evaluate the enrichment of common genetic variants from genome-wide association study (GWAS) signals to partition trait heritability within functional categories represented by our DMRs, ATAC peaks, and DAPs^26^. LDSC estimates the proportion of genome-wide single nucleotide polymorphism (SNP)-based heritability that can be attributed to SNPs within a given category by a regression model that combines GWAS summary statistics with estimates of linkage disequilibrium from an ancestry-matched reference panel. We ran LDSC (v1.0.0; https://github.com/bulik/ldsc) to estimate the proportion of genome-wide SNP-based heritability across 25 traits within 7 categories defined from our set of DMRs and DAPs. Each of these 7 categories was added one at a time to a ‘full baseline model’ that included 53 functional categories (24 main annotations, 500 bp windows around of each of the 24 main annotations, and 100 bp windows around 5 sets of ChIP-seq peaks) that capture a broad set of genomic annotations, as previously described^26^.

The 7 categories were:

1. Neuronal DMRs (Supplementary Table 5)
2. Neuronal DAPs (Supplementary Table 16)
3. Neuronal blocks (Supplementary Table 10)
4. Neuronal non-DAPs^1^ (Supplementary Table 16)
5. Neuron vs. glia DMRs (Supplementary Table 11)
6. Neuron vs. glia DAPs (Supplementary Table 22)
7. Neuron vs. glia blocks (Supplementary Table 12)

The 25 traits and their summary statistics were:

1. ADHD^27^ (https://www.med.unc.edu/pgc/files/resultfiles/pgc.adhd.2012-10.zip)
2. Age at menarche^28^ (http://www.reprogen.org/Menarche_Nature2014_GWASMetaResults_17122014.zip)
3. Alzheimer’s disease^29^ (IGAP_summary_statistics.zip downloaded from http://web.pasteur-lille.fr/en/recherche/u744/igap/igap_download.php)
4. Autism spectrum disorder (Autism Spectrum Disorder Working Group of the Psychiatry Genomics Consortium. Dataset: PGC-ASD summary statistics from a meta-analysis of 5,305 ASD-diagnosed cases and 5,305 pseudocontrols of European descent (based on similarity to CEPH reference genotypes, March 2015; https://www.med.unc.edu/pgc/files/resultfiles/pgcasdeuro.gz)
5. Bipolar disorder^30^ (https://www.med.unc.edu/pgc/files/resultfiles/pgc.bip.2012-04.zip)
6. Cognitive performance^31^ (http://ssgac.org/documents/CHIC_Summary_Benyamin2014.txt.gz)
7. Depressive symptoms^32^ (http://ssgac.org/documents/DS_Full.txt.gz)
8. Educational attainment^33^ (http://ssgac.org/documents/SSGAC_Rietveld2013.zip)
9. Neuroticism^32^ (http://ssgac.org/documents/Neuroticism_Full.txt.gz)
10. PGC cross-disorder^27^ (https://www.med.unc.edu/pgc/files/resultfiles/pgc.cross.full.2013-03.zip)
11. Schizophrenia^34^ (https://www.med.unc.edu/pgc/files/resultfiles/scz2.snp.results.txt.gz)
12. Subjective well-being^32^ (http://ssgac.org/documents/SWB_Full.txt.gz)
13. Age of smoking onset^35^ (https://www.med.unc.edu/pgc/files/resultfiles/tag.logonset.tbl.gz)
14. Cigarettes smoked per day^35^ (https://www.med.unc.edu/pgc/files/resultfiles/tag.cpd.tbl.gz)
15. Ever smoked^35^ (https://www.med.unc.edu/pgc/files/resultfiles/tag.evrsmk.tbl.gz)
16. Former vs. current smoker^35^ (https://www.med.unc.edu/pgc/files/resultfiles/tag.former.tbl.gz)
17. HDL^36^ (http://archive.broadinstitute.org/mpg/pubs/lipids2010/HDL_ONEEur.tbl.sorted.gz)
18. Height^37^ (http://portals.broadinstitute.org/collaboration/giant/images/4/49/GIANT_HEIGHT_LangoAllen2010_publicrelease_HapMapCeuFreq.txt.gz)
19. LDL^36^ (http://archive.broadinstitute.org/mpg/pubs/lipids2010/LDL_ONEEur.tbl.sorted.gz)
20. Triglycerides (TG)^36^ (http://archive.broadinstitute.org/mpg/pubs/lipids2010/TG_ONE_Eur.tbl.sorted.gz)
21. BMI^38^ (http://portals.broadinstitute.org/collaboration/giant/images/b/b7/GIANT_BMI_Speliotes2010_publicrelease_HapMapCeuFreq.txt.gz)
22. Coronary artery disease^35^ (http://www.cardiogramplusc4d.org/media/cardiogramplusc4d-consortium/data-downloads/cardiogram_gwas_results.zip)
23. Fasting glucose^39^ (ftp://ftp.sanger.ac.uk/pub/magic/MAGIC_Manning_et_al_FastingGlucose_MainEffect.txt.gz)
24. Rheumatoid arthritis^40^ (http://plaza.umin.ac.jp/yokada/datasource/files/GWASMetaResults/RA_GWASmeta_European_v2.txt.gz)
25. Type-2 diabetes^41^ (DIAGRAMv3.2012DEC17.zip, corresponding to ‘Stage 1 GWAS: Summary Statistics’, downloaded from http://www.diagram-consortium.org/downloads.html)

For each of the 7 × 25 = 175 category/trait combinations, we used LDSC to estimate the category’s enrichment, the associated standard error, and the associated P-value. Enrichment is calculated as the proportion of SNP heritability accounted for by each category divided by the proportion of total SNPs within the category; its standard error is assessed using a block jack-knife procedure. Using the enrichment and its standard error, LDSC forms a Z-score and associated P-value, which we subsequently adjusted for multiple testing using the method of Benjamini and Hochberg^42^.

The baseline categories were included in all 7 fits per trait. There was some slight variation in the estimates for these baseline categories across the 7 fits. To summarize each baseline-category/trait combination, we selected the enrichment and standard error associated with the minimum P-value across the 7 fits. For clarity of presentation, we only show results for two of the baseline categories: ‘Conserved’ (the most enriched category across a broad range of traits in a previous analysis^26^) and ‘FANTOM5 enhancer’; these categories serve as control regions for comparison with our DMRs and DAPs.

### Enrichment of de novo mutations within DMRs and DAPs

We downloaded a list of de novo mutations (DNMs) from 192 individuals with autism (Supplementary Table 443), specifically, containing 10,387 single nucleotide variants. We also downloaded a list of de novo mutations from 258 control individuals^44^, specifically the GoNL_DNMs.txt file from http://www.nlgenome.nl/?page_id=9 containing 11,020 single nucleotide variants.

For each set of DNMs, we constructed a 2×2 contingency table cross-classifying whether each base in the autosomal genome was a DNM and within a neuronal DMR. We then estimated the log odds ratio for the enrichment of DNMs within DMRs as in section ‘**Enrichment of DMRs, ATAC peaks, and DAPs in genomic features**’. We repeated the analysis for neuronal DAPs.

We also used a simple simulation to check whether:

1. The DMRs/DAPs contained more DNMs than we would expect by chance
2. The proportion of individuals with at least one DNM in a DMR/DAP was larger than we would expect by chance.

Briefly, we sampled 10,000 null sets of regions from the autosomal genome. Each set of null regions had as many regions as there were DMRs/DAPs and with the same width distribution as the DMRs/DAPs. For each set of null regions, we tabulated the number of DNMs in each set overlapping the null regions and the number of individuals with at least one DNM in a null region. We then compared the observed counts to the counts from the simulated null regions to derive empirical P-values for each hypothesis. This analysis was performed for both the set of DNMs from individuals with autism and the set of DNMs from control individuals.

### Transcription Factor Motif Enrichment

We used the Haystack (v0.4)^45^ `haystack_motifs` module to scan for vertebrate JASPAR (2016)^46^ transcription factor binding motifs enriched in our datasets. We separated the hypo-, hypermethylated DMR and hypo-, hyper-DAPs identified between NAcc and BA9 in NeuN^+^ cells into DMRs or DAPs that overlapped a promoter and those that did not, resulting in eight lists that were input into Haystack as BED files. All autosomal promoters were input as background for the DMRs or DAPs that overlapped promoters while the entire autosomal hg19 genome excluding the promoters was input as background for the DMRs or DAPs not overlapping promoters. Haystack selects a random, CG content-matched subset of the input background to use for enrichment calculations.

### Gene Ontology and KEGG Annotation

We utilized the Genomic Regions Enrichment of Annotations Tool (GREAT; version 3.0.0)^47^ to assess nearest gene enrichment for distinct sets of DMRs and DAPs we identified. We used the hg19 assembly, reduced the default input parameter for max extension to 100 kb, and kept all other default parameters the same (settings available at the GREAT website: http://great.stanford.edu/). Gene Ontology (GO) terms returned must be significant by both the binomial and hypergeometric tests using the multiple hypothesis correction false discovery rate (FDR) ≤ 0.05 whose binomial fold enrichment is at least 2.0. If no GO terms were enriched, we reported MSigDB Pathway results if present.

EnrichR was used to perform gene ontology (GO) and KEGG pathway analyses using lists of gene symbols as input. Gene lists were generated by matching GENCODE gene IDs to gene symbols (“external_gene_id”) using biomaRt^48,49^.

### MEF2C ChIP from ENCODE

ChIP data generated by ENCODE for the MEF2C transcription factor in lymphoblastoid cells (GM12878) was downloaded using AnnotationHub on 8/15/2016 (AnnotationHub ID: AH22613; URL: http://hgdownload.cse.ucsc.edu/goldenpath/hg19/encodeDCC/wgEncodeAwgTfbsUniform/wgEncodeAwgTfbsHaibGm12878Mef2csc13268V0416101UniPk.narrowPeak.gz).

### Software

All statistical analyses were performed using R^24^ (v3.3.x) and made use of packages contributed to the Bioconductor project^50,51^. In addition to those R/Bioconductor packages specifically referenced in the above, we made use of several other packages in preparing results for the manuscript:

- AnnotationHub (v1.36.2)
- biomaRt^48,49^ (v2.30.0)
- GenomicAlignments^16^ (v1.10.0)
- GenomicFeatures^16^ (v1.26.2)
- GenomicRanges^16^ (v1.26.2)
- ggplot2^52^ (v2.2.1)
- Hmisc (v4.0-2)
- Matrix (v1.2-8)
- rtracklayer^53^ (v1.34.1)
- SummarizedExperiment (v1.4.0)

## Footnotes

^1^‘Non-DAPs (Neuron)’ were all those ATAC peaks not called as ‘DAPs (Neuron)’, i.e. adjusted P-value > 0.05 and/or |logFC| < 1 in comparison of NAcc and BA9 in NeuN^+^ cells.

